# Gene-Corrected Basal Cells Restore CFTR In Vitro; Transplants Regenerate Epithelium in a Preclinical Sinus Model

**DOI:** 10.1101/2025.07.21.666023

**Authors:** Katelin M. Allan, Shaoyan Zhang, Sharon L. Wong, Alexander Capraro, Alexandra McCarron, Daniel Skinner, Egi Kardia, Ye Zheng, Ling Zhong, Chai-Ann Ng, Jessica L. Bell, Nigel Farrow, Bala Umashankar, Catherine Banks, Cristan Herbert, Kristopher A. Kilian, Elvis Pandzic, Orazio Vittorio, Jamie I. Vandenberg, John G. Lock, Adam Jaffe, Martin Donnelley, David Parsons, Bradford A. Woodworth, Do-Yeon Cho, Shafagh A. Waters

**Author notes:** Correspondence should be addressed to S.A.W, Wallace Wurth Building UNSW Sydney 2052. **Take home message** *In vitro CFTR* gene addition restores CFTR activity and mucociliary function in airway basal cells across genotypes, including nonsense mutations. *In vivo*, transplanted basal cells regenerate functional airway epithelium in a rabbit sinus model.

## Abstract

**Background:** Cystic fibrosis (CF) is caused by mutations in the *CFTR* gene, leading to epithelial dysfunction and progressive lung disease. Although CFTR modulators have transformed care, ∼10% of people with CF remain without effective therapy. Durable, mutation-agnostic approaches are urgently needed.

**Method:** We used a lentiviral (LV) vector to deliver wild-type *CFTR* to airway basal cells derived from 13 paediatric CF participants with a range of genotypes. Transduced cells were assessed for transgene expression, epithelial differentiation, and CFTR function using air-liquid interface (ALI) cultures. Separately, to evaluate regenerative capacity *in vivo*, LV^GFP^-transduced rabbit airway basal cells were transplanted into the denuded nasal septum of healthy New Zealand white rabbits using a biocompatible scaffold.

**Results:** Transduced basal cells retained multilineage differentiation capacity, forming well-organized, pseudostratified epithelium with intact barrier function and ciliary activity. CFTR channel activity was restored to levels comparable to or exceeding those achieved with elexacaftor/tezacaftor/ivacaftor (ETI), including in individuals with nonsense mutations. Combined *CFTR* transduction plus ETI treatment showed additive benefit. *In vivo*, transplanted rabbit basal cells engrafted and differentiated to regenerate a mucociliary epithelium, with improved nasal potential difference and mucociliary clearance compared to scaffold-only controls.

**Conclusion:** Our study demonstrates that LV-mediated *CFTR* gene addition restores CFTR function *in vitro* across genotypes and supports epithelial regeneration in a clinically relevant animal airway model. This two-part platform offers a scalable path toward cell therapies for all people with CF and may have broader applications in upper airway epithelial repair.

## Introduction

Cystic Fibrosis (CF) is caused by pathogenic variants of the *CF transmembrane conductance regulator* (*CFTR*) gene. Among the more than 2,121 *CFTR* variants identified to date [1], many result in loss of CFTR protein function and lead to multisystem disease. The CFTR protein functions as an epithelial anion channel, regulating chloride (Cl⁻) transport at the apical surface of epithelial tissues [2]. Although small molecule CFTR modulators, such as elexacaftor/tezacaftor/ivacaftor (ETI), are now available for individuals carrying at least one responsive *CFTR* mutation, clinical benefit is variable, and approximately 10% of people with CF (pwCF) remain without effective modulator options [3]. Moreover, modulator-based correction is transient and genotype-restricted. These limitations highlight the ongoing need for durable, mutation-agnostic therapeutic strategies to restore CFTR function across all genotypes (reviewed in [4]).

Gene therapy targeting airway basal cells represents a promising strategy for durable epithelial correction. Basal cells serve as tissue-resident progenitors capable of regenerating the full spectrum of specialised epithelial lineages, including ciliated, secretory and ionocyte cells [5, 6]. Lentiviral (LV) vectors are well-suited for this task, since they accommodate the full-length *CFTR* cDNA transgene and stably integrate into the host genome [7]. However, direct *in vivo* delivery to basal cells remains challenging due to their subapical location beneath intact airway epithelium [8]. Conditioning strategies, such as lysophosphatidylcholine (LPC) administration or mechanical epithelial disruption, have been used to transiently expose basal cells and enhance transduction efficiency [9]. While long-term transgene expression has been achieved in animal models (up to 18 months), overall *in vivo* transduction efficiency remains suboptimal [10, 11].

*Ex vivo* gene correction followed by transplantation of airway basal cells offers an alternative strategy to circumvent these *in vivo* delivery barriers [4]. Prior studies have demonstrated that CFTR-corrected basal cells from the bronchi, sinuses, and trachea can differentiate into polarized epithelium with improved ion transport *in vitro* [12–16] and retain CFTR activity when cultured on a bioscaffold *ex vivo* [14]. While the ability of basal cells to engraft and regenerate mucociliary architecture *in vivo* has been reported [17], restoration of function remains incompletely defined.

To date, most *in vivo* airway cell transplantation studies have been conducted in non-CF contexts and human clinical trials remain limited to the lower airways, with evaluation often focused on gross histology and spirometry [18–22]. A robust, multiparametric framework is needed to fully assess the therapeutic potential of *CFTR*-corrected basal cell therapies. The upper airways, particularly the paranasal sinuses, offer a clinically relevant site for early phase testing of airway regenerative approaches. In CF, the sinuses serve as persistent bacterial reservoirs that can perpetuate lower airway infection, including post-lung transplantation where re-colonization from the upper airway remains a major clinical challenge [23]. Their epithelial composition closely resembles that of the lower airways, and their surgical accessibility makes them ideal for localized delivery and engraftment studies [24].

In this study (**Figure 1**), we aimed to assess whether LV-mediated CFTR correction of airway basal cells could restore epithelial structure and function *in vitro* and whether corrected cells retain engraftment potential *in vivo*. For *in vitro* analysis, we transduced airway basal cells from 13 paediatric CF participants with an LV-*CFTR* vector (LV*^CFTR^*) and assessed differentiation at air-liquid interface (ALI). We evaluated epithelial morphology, barrier integrity, global proteomic profiling, ciliary function and CFTR activity – including ETI responsiveness – relative to untransduced (naive) CF epithelium and non-CF epithelium (**Supplementary Table 1**).

**Figure 1.**
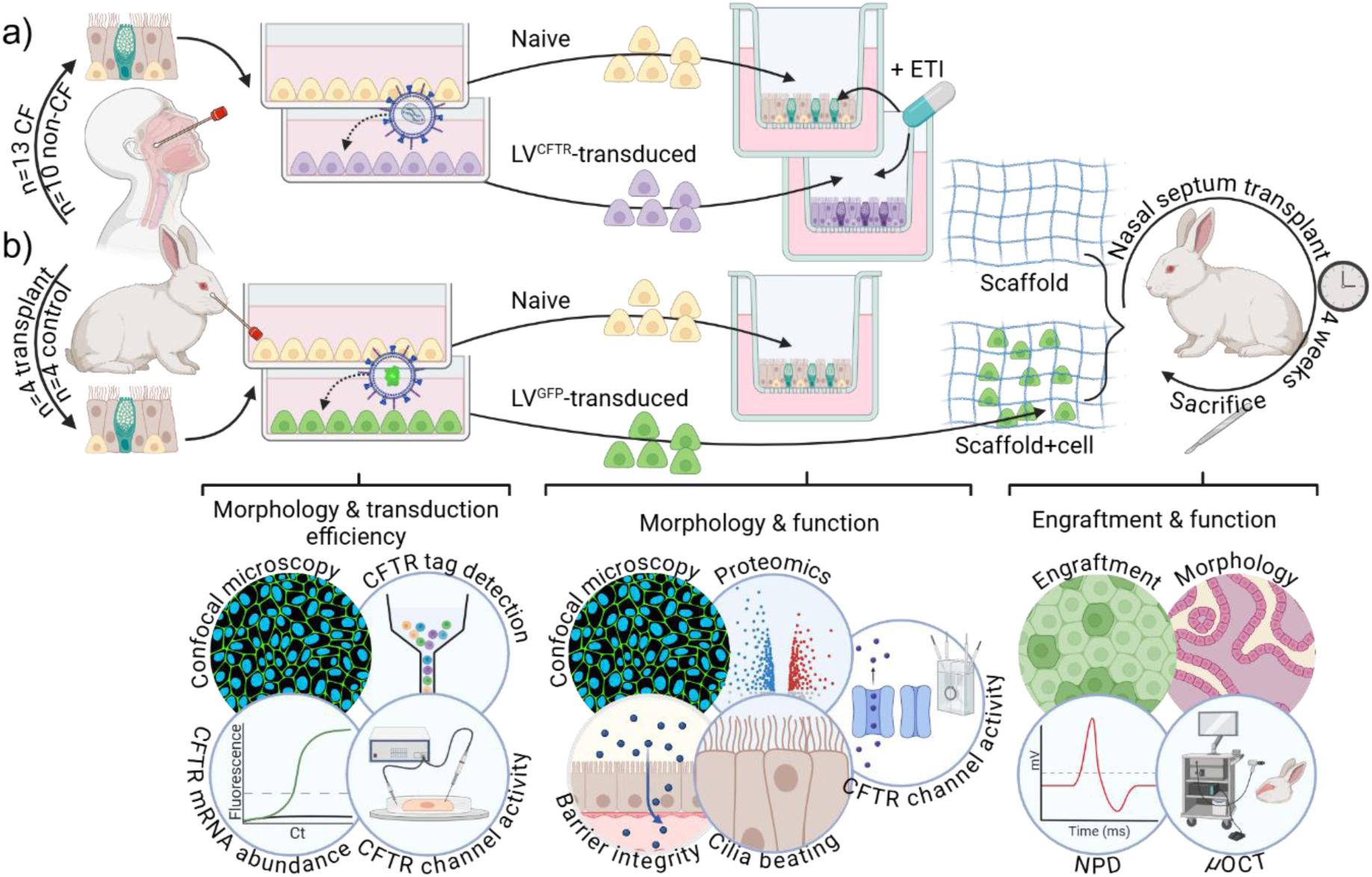
Schematic of study design. **a)** Airway mucosa were collected via nasal brushings from paediatric CF participants (n=13) and non-CF controls (n=10). Basal cells were expanded in culture and transduced with a lentiviral vector (LV) carrying the wild-type *CFTR* transgene (LV*^CFTR^*). Morphology was assessed post-transduction, and transduction efficiency was quantified by multiple methods (flow cytometry, qPCR and automated single-cell patch clamp analysis). Naive (untransduced) and transduced basal cells were differentiated at air-liquid interface (ALI) to form pseudostratified airway epithelium. Structural and functional assessments including epithelial morphology, barrier integrity, global proteomic profiling, ciliary beat frequency and CFTR channel function were performed. CFTR modulator treatment (elexacaftor/tezacaftor/ivacaftor; ETI) was applied post-differentiation to naive- and transduced-derived epithelium to assess potential additive effects on CFTR activity. **b)** Rabbit airway basal cells were collected, expanded and either differentiated at ALI to validate airway epithelium morphology and CFTR function *in vitro*, or transduced with LV-eGFP reporter and embedded in an FDA-approved bioscaffold for nasal septum transplantation (n=4 transplant; n=4 control). Engraftment was assessed after four weeks by GFP expression, with epithelial morphology and mucociliary function evaluated by histology, *ex vivo* micro-optical coherence tomography (µOCT), and nasal potential difference (NPD) performed prior to the animals being humanely killed.

For *in vivo* testing, we selected the rabbit paranasal sinus as a prototypical airway regeneration model due to its anatomical and histological similarity to the human sinus epithelium, featuring pseudostratified columnar architecture, robust mucociliary clearance, and surgical accessibility [25]. We transplanted LV^GFP^-transduced rabbit airway basal cells embedded in an FDA-approved bioscaffold (Myriad Matrix) into the mechanically disrupted nasal septum of healthy rabbits. Engraftment, epithelial regeneration, and mucociliary function were assessed four weeks post-transplantation.

## Methods

Human ethics approval was granted by the Sydney Children’s Hospital Network (HREC/16/SCHN/120); animal studies were approved by the University of Alabama at Birmingham Institutional Animal Care and Use Committee (IACUC-22329).

Detailed materials and methods are provided in the **Supplementary Material**. Briefly, primary airway epithelial cells were collected from CF (n=13; W1282X/W1282X, G542X/F508del, Q493X/F508del, F508del/F508del) and non-CF (n=10) participants (**Supplementary Table 2**), and expanded as basal cell monolayers using the conditionally reprogrammed cell method [26]. Basal cells were transduced with a lentiviral vector (LV) carrying wild-type *CFTR*. *CFTR* transduction efficiency was assessed via flow cytometry, qPCR, and automated single-cell patch clamp. Naive and transduced basal cells were differentiated at air-liquid interface (ALI) until maturity [26], and evaluated for epithelial composition, barrier integrity, ciliary beat frequency, global proteomics, and CFTR function [27]. ETI was tested both alone and in combination with transduced-derived epithelium.

Rabbit nasal epithelial cells were similarly isolated, expanded, and differentiated at ALI, with immunofluorescence confirming normal epithelial composition and Ussing chamber electrophysiology verifying CFTR activity. Bioscaffolds – either cell-free (n=4) or seeded with LV^GFP^-transduced rabbit basal cells (n=4) – were transplanted onto the mechanically denuded nasal septum of New Zealand white rabbits. Grafts were assessed after four weeks by histology, immunofluorescence, micro-optical coherence tomography (µOCT), and nasal potential difference (NPD) measurements [28].

## Results

### Morphology of airway basal cells is preserved post LV^CFTR^ transduction

Primary basal cell monolayers were established from paediatric CF human nasal epithelial cells (hNECs) carrying W1282X/W1282X, G542X/F508del, or F508del/F508del genotypes (**Supplementary Table 1**). Monolayers were transduced with a lentiviral (LV) vector encoding V5-tagged wild-type *CFTR* (LV*^CFTR^*). V5 expression was detected exclusively in *CFTR-*transduced (Td*^CFTR^*) basal cells, which retained a stable morphology (e.g., cell size and shape) compared to untransduced (naive) controls (**Supplementary Figure 1a-b**).

At a multiplicity of infection (MOI) of 10, *CFTR* transduction efficiency showed inter-participant variability but was quantifiable by multiple orthogonal methods: flow cytometry (14.0 ± 4.3 %), qPCR (635.6 ± 358.3 2^-ΔΔCt^) and automated patch clamping (14.3 ± 5.2%, −192.5 ± 15.4 pA/pF) (**Supplementary Figure 1c-g, 2, 3a-b**). Importantly, endogenous F508del *CFTR* mRNA levels remained unchanged post-transduction (**Supplementary Figure 1h**), indicating that LV*^CFTR^* does not alter mutant gene expression. Repeating the transduction procedure did not enhance efficiency (**Supplementary Figure 1i**).

To explore whether higher doses might improve efficiency, a follow-up experiment in a CF subset (n=6) demonstrated that increasing the MOI to 20 significantly improved transduction efficiency by 1.6-fold (14.3 ± 2.1% vs. 9.1 ± 2.8; *p*<0.01; **Supplementary Figure 4a-b**). V5-positive Td*^CFTR^*basal cell frequency showed a moderate correlation with *CFTR* transgene mRNA expression levels (ρ = 0.67; **Supplementary Figure 5a**), and a strong positive correlation with the number of cells expressing CFTR current (ρ = 0.96, *p* < 0.01; **Supplementary Figure 5b**). In contrast, CFTR activity correlated only weakly with transgene mRNA levels (ρ = 0.43; **Supplementary Figure 5c**), underscoring the importance of functional protein expression over transcript abundance.

### Transduced^CFTR^ basal cells retain their capacity for multilineage epithelial differentiation

To assess whether LV*^CFTR^* transduction impacts epithelial differentiation, we compared naive-derived and Td*^CFTR^*-derived epithelium following air-liquid interface (ALI) culture of primary CF airway basal cells. Td*^CFTR^*-derived epithelium successfully differentiated into a well-organised, pseudostratified epithelium with apical-basal polarity resembling that of naive-derived mucociliary epithelium (Figure 2a-c, **Supplementary Figure 6**).

**Figure 2.**
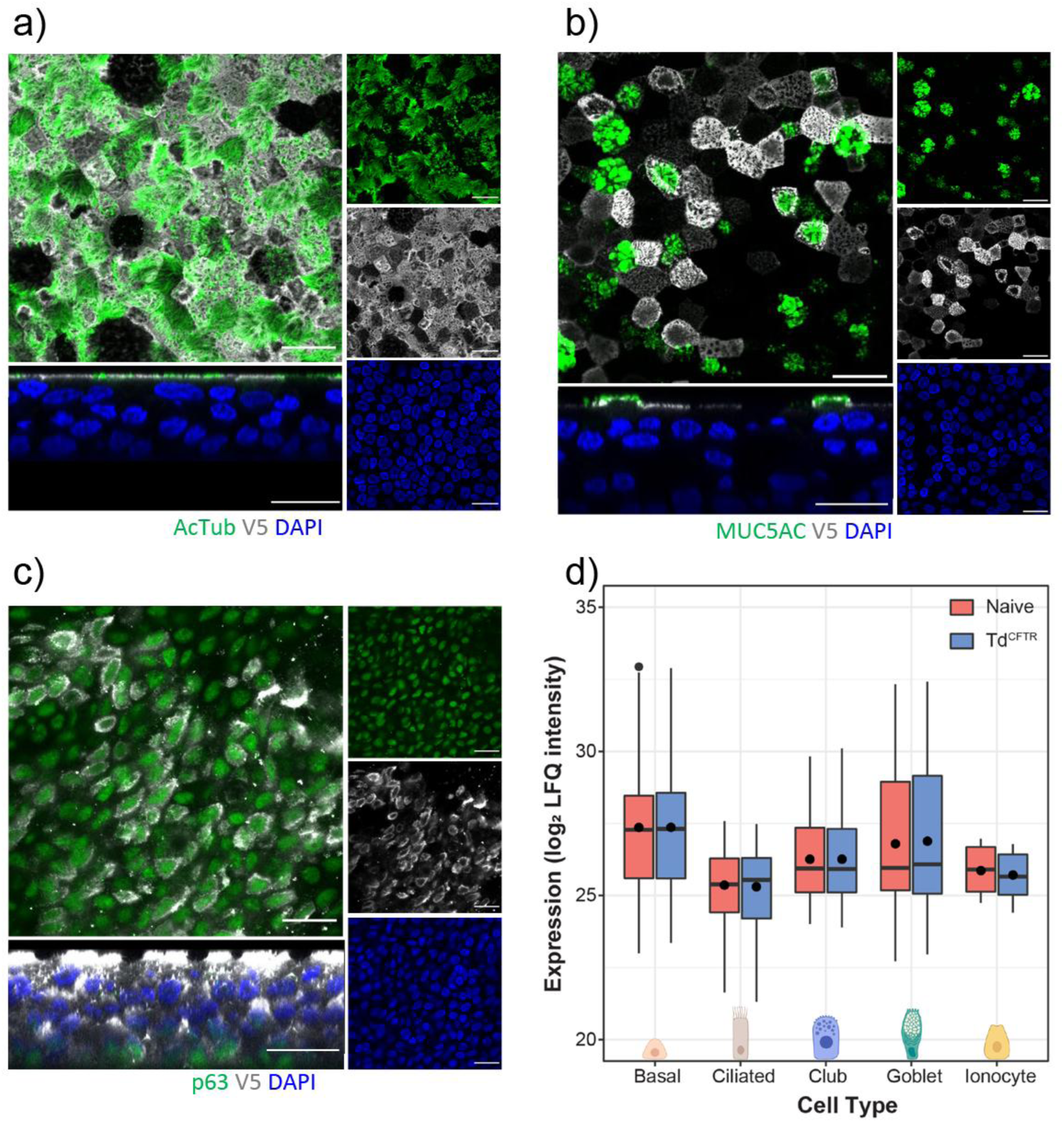
Epithelial morphology and cellular composition of transduced basal cells differentiated at air-liquid interface. **a-c)** Immunofluorescence staining of air-liquid interface cultures showing V5-tagged *CFTR* (grey) co-localised with **a)** ciliated cells (acetylated tubulin, AcTub, green), **b)** secretory goblet cells (MUC5AC, green) and **c)** basal cells (p63, green). Bottom left panels show z-stacks. Remaining panels show top-down view. Images captured using a 63×/1.4 oil immersion objective. Scale bars = 20 μm. Additional images are provided in Supplementary Figure 6. **d)** Expression levels of major airway epithelial cell types in naive (n=10) and transduced (Td*^CFTR^*)-derived (n=10) CF epithelium, measured by average log_2_ label-free quantification (LFQ) of mass spectrometry data. Boxplots show the median, first and third quartiles, with whiskers representing the largest and smallest values no greater than 1.5× interquartile range from the box. Statistical significance was determined by Wilcoxon rank sum test.

Immunofluorescence revealed appropriate spatial localisation of key cell-type markers: ciliated (acetylated tubulin) and goblet (MUC5AC) cell markers were confined to the apical surface, while basal cell marker (p63) localised to the basal membrane. V5-tagged CFTR was detectable across all epithelial layers, but was not expressed uniformly across all cells, indicating heterogenous transduction of the original basal cells. Co-localisation with ciliated, secretory, and basal markers confirmed that CFTR expression persisted throughout differentiated cell types (**Figure 2a-c**). Global proteomics profiling further supported this, showing no significant difference in epithelial cell-type marker expression between Td*^CFTR^*- and naive-derived epithelium (**Figure 2d**).

Tight junction protein expression (OCLN, ZO-1, ZO-2, ZO-3) was maintained across both groups (**Figure 3a**), and phalloidin staining confirmed apical actin distribution consistent with a well-polarised epithelial barrier (**Figure 3b**). Functional epithelial integrity was supported by comparable transepithelial electrical resistance (TEER) values (naive: 509.3 ± 84.0 Ω.cm^2^, Td*^CFTR^*: 549.2 ± 71.9 Ω.cm^2^; **Figure 3c**). Cilia beat frequency (CBF) remained within physiological range and showed no significant difference between naive and Td*^CFTR^*-derived epithelium (8.74 ± 0.48 vs. 9.16 ± 0.39; **Figure 3d**). Notably, the proportion of V5-positive basal cells moderately correlated with CBF in the resultant epithelium (ρ = 0.70, *p* < 0.05; **Supplementary Figure 5d**), suggesting that CFTR expression may enhance motile cilia function.

**Figure 3.**
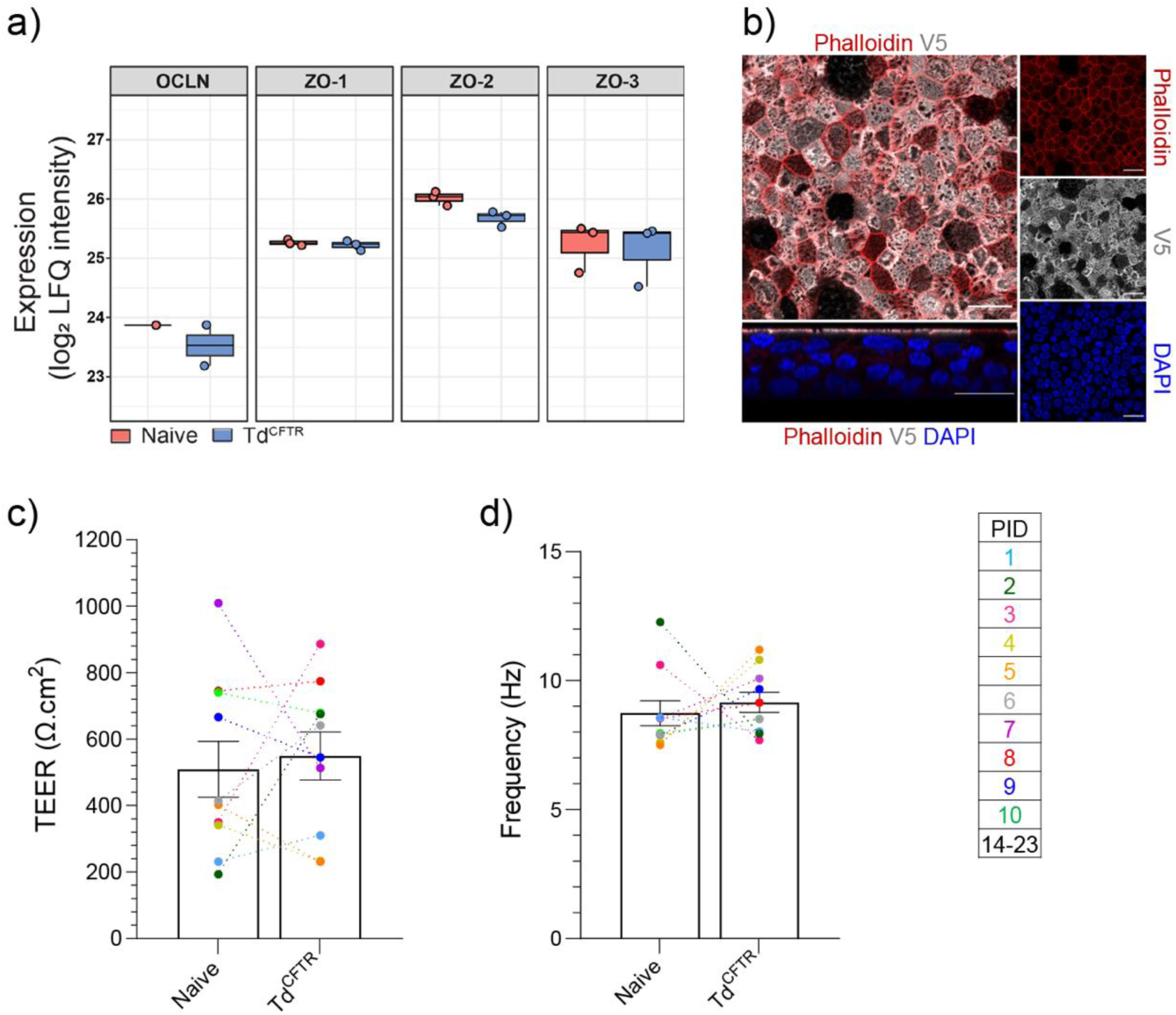
Epithelial barrier integrity and ciliary function in transduced-derived epithelium. **a)** Expression of tight junction proteins (OCLN, ZO-1, ZO-2 and ZO-3) in naive (n=10) and transduced-derived (n=10) CF epithelium, measured by average log_2_ label-free quantification (LFQ) of mass spectrometry data. Boxplots show the median, first and third quartiles, with whiskers representing the largest and smallest values no greater than 1.5× interquartile range from the box. Statistical significance was determined by Wilcoxon rank sum test. **b)** Immunofluorescence staining of actin (phalloidin, red) localised to intercellular junctions in epithelial cells expressing V5-tagged *CFTR* (grey). Images captured with a 63×/1.4 oil immersion objective. Scale bars = 20 μm. **c)** Dot plot of mean transepithelial electrical resistance (TEER_;_ Ω.cm^2^) and **d)** cilia beat frequency (Hz) in naive and transduced-derived epithelium of CF participants (n=10). Data are presented as mean ± SEM, with each dot representing the average of n=3 replicate ALI cultures per participant. PID=Participant ID. hNEC=human nasal epithelial cell. Statistical significance was determined using paired t tests.

### Transduced^CFTR^-derived epithelium maintains cellular identity and secretory landscape

To evaluate whether LV*^CFTR^* transduction alters epithelial proteomic profiles, we performed global proteomic analysis of the two groups. Only a small fraction of proteins was differentially abundant in Td*^CFTR^*-derived epithelium compared to naive-derived epithelium. Specifically, 1.8% of intracellular (cell lysate) and 3.8% of secreted (apical secretome) proteins (**Figure 4a**). Most changes were modest and primarily enriched for metabolic pathways, including inositol phosphate metabolism, which has been previously associated with LV transduction [29] (**Figure 4b, Supplementary Table 3**). Importantly, similar minimal proteomic shifts were observed in LV^eGFP^-derived epithelium, reinforcing that changes were vector-related rather than transgene-specific (**Supplementary Figure 7a-b, Supplementary Table 3**).

**Figure 4.**
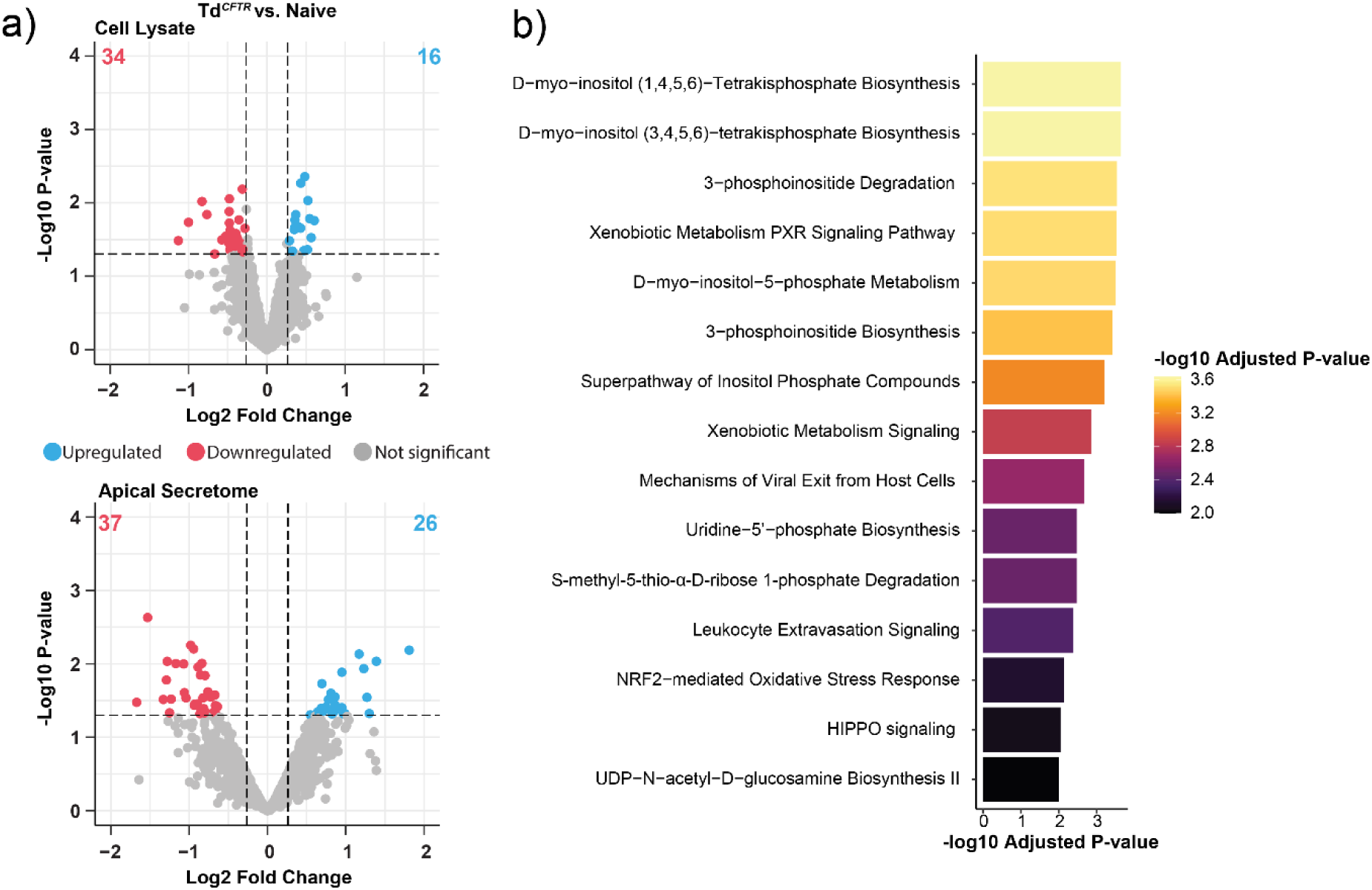
Global proteomic profiling of transduced-derived airway epithelium. **a)** Volcano plots showing differentially abundant proteins in the cell lysate (top, n=10) and apical secretome (bottom, n=3) of transduced (Td*^CFTR^*)-derived epithelium compared to naive-derived epithelium. Each dot represents an individual protein. Upregulated proteins are shown in blue, downregulated in red, and non-significant proteins in grey. Total numbers of differentially abundant proteins are annotated in colour. Dotted lines indicate statistical thresholds (*p*-value < 0.05; fold change > |1.2|). Statistical analysis was performed using the DEP R Package (see methods). **b)** Top canonical pathways enriched among differentially abundant proteins in cell lysate, identified via Ingenuity Pathway Analysis (IPA). Bar colour reflects −log_10_ adjusted *p*-value. See Supplementary Table 3 for full pathway lists.

### CFTR channel activity is restored in Transduced^CFTR^-derived epithelium and further enhanced by ETI treatment

To evaluate functional CFTR restoration, Ussing chamber assays were performed on differentiated hNECS from 13 paediatric CF participants spanning a range of CFTR genotypes: seven with Class II (F508del/F508del: PID 7-13) and six with at least one Class I mutation (W1282X/W1282X, G542X/F508del, Q493X/F508del; PID 1-6).

Naive-derived epithelium from all CF participants showed negligible CFTR activity (<3.00 μA/cm^2^; **Figure 5a**). In contrast, Td*^CFTR^*-derived epithelium exhibited significantly increased CFTR activity (−2.9 to −50.8 μA/cm^2^, *p*<0.05; **Figure 5a**), which correlated strongly with transduction efficiency (ρ = - 0.92, *p* < 0.0001; **Figure 5b**) and the number of cells expressing CFTR current (ρ = −0.79, *p* < 0.05; **Supplementary Figure 8a**), and moderately with *CFTR* transgene expression (ρ = −0.57, *p* = ns; **Supplementary Figure 8b**). CFTR activity levels varied between individuals with higher transduction associated with greater functional improvement.

**Figure 5.**
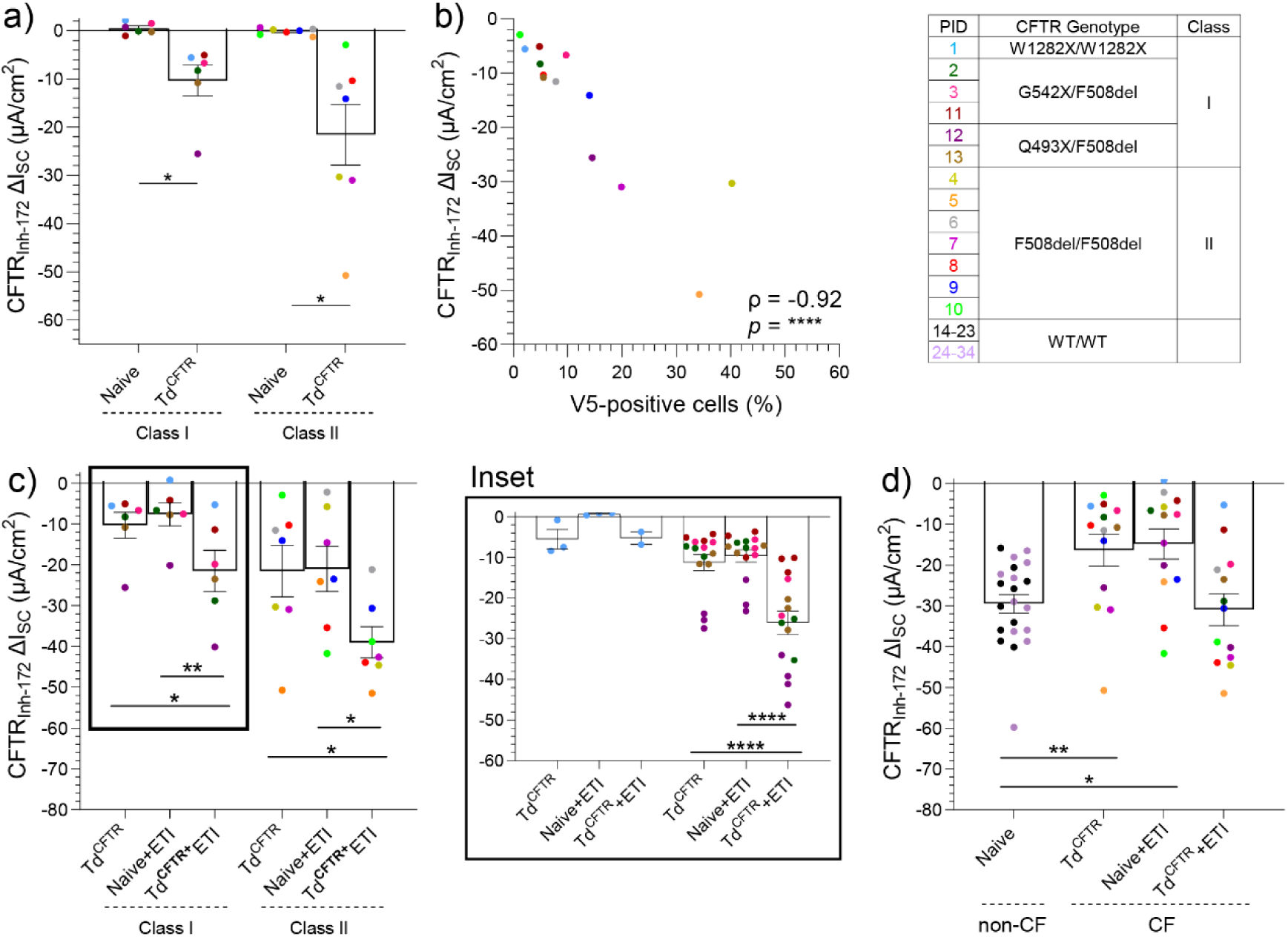
Restoration of CFTR channel activity in Td*^CFTR^*-derived epithelium, with and without ETI treatment. **a)** CFTR-mediated chloride currents (CFTR-Inh_172_) in naive versus Td*^CFTR^*-derived epithelium from 13 CF participants spanning Class I and Class II *CFTR* genotypes. **b)** Spearman’s correlation between V5-positive basal cell frequency (*CFTR* transgene) and CFTR-Inh_172_ current (μA/cm^2^). **c)** Comparison of CFTR activity across three conditions: naive-derived plus ETI-treated (naive+ETI), Td*^CFTR^*, and Td*^CFTR^*-derived plus ETI-treated (Td*^CFTR^*+ETI) epithelium. Inset: Individual replicate values are shown for each donor with Class I genotypes (W1282X/W1282X, n=1; G542X/F508del, n=3; Q493X/F508del, n=2), with n=2-4 replicate ALI cultures per participant. **d)** Summary of all treatment conditions in CF epithelium (n=10) compared to CFTR activity in non-CF epithelium from this study (n=10; black) and our prior cohort (n=11; purple). Each point represents the mean from three replicate ALI cultures (unless otherwise noted). Error bars = SEM. Statistics: paired t test (a), Spearman’s correlation (b), One-way ANOVA with the Geisser-Greenhouse correction and Tukey’s multiple comparison test (c) or Brown-Forsythe or Welch ANOVA tests with Dunnet’s T3 multiple comparison test (inset, d). \**p*<0.05, \*\*\**p*<0.001, \*\*\*\**p*<0.0001. PID=Participant ID. ETI=elexacaftor/tezacaftor/ivacaftor.

In a subset (n=6), increasing the MOI from 10 to 20 significantly enhanced CFTR activity by 1.6-fold, without affecting TEER or CBF (**Supplementary Figure 4c-e**). This increase remained strongly correlated with transduction efficiency (ρ = −0.83, *p* < 0.01; **Supplementary Figure 4f**). CFTR activity was transgene-specific: LV^eGFP^-derived epithelium exhibited negligible CFTR activity, comparable to naive-derived epithelium and significantly lower than Td*^CFTR^*-derived epithelium (*p*<0.001; **Supplementary Figure 9**, **Supplementary Table 4**). Comparable trends were observed in Td*^CFTR^*-derived bronchial epithelium, though direct nasal-to-bronchial comparisons were limited by variable transduction efficiencies (n=3; **Supplementary Figure 10a-c**).

To benchmark against clinical therapy, we treated naive-derived epithelium with ETI. As expected, the W1282X/W1282X participant (PID 1) showed no response. In contrast, G542X/F508del and Q493X/F508del participants exhibited significant increases in CFTR activity (Δ: −9.8 μA/cm^2^, *p*<0.0001; **Supplementary** Figure 11 **inset**). F508del/F508del participants also responded significantly to ETI (Δ: −20.9 μA/cm^2^, *p*<0.01; **Supplementary Figure 11**).

Direct comparisons showed that Td*^CFTR^*-derived epithelium restored CFTR activity to levels similar to or exceeding those achieved by ETI-treated naive-derived epithelium (**Figure 5c**). In Td*^CFTR^*-derived epithelium from class I mutation carriers G542X/F508del and Q493X/F508del, CFTR activity levels were similar to those achieved by ETI in naive-derived epithelium (**Figure 5c**). In case of PID1, an individual homozygous for the W1282X nonsense mutation, a known non-responder to ETI, CFTR activity in naive-derived epithelium remained negligible even after ETI treatment (naive-ETI: 0.73 ± 0.17 μA/cm^2^; **Figure 5c inset**). However, Td*^CFTR^*-derived epithelium from this individual showed a CFTR activity increase (Δ: −6.3 μA/cm^2^), despite a transduction efficiency of only 2.1% (**Supplementary Figure 12**). In the epithelium derived from F508del/F508del participants, responses varied: some achieved significantly higher CFTR activity with Td*^CFTR^*, others with ETI (**Supplementary Figure 12**).

To determine potential synergy, we applied ETI to Td*^CFTR^*-derived epithelium. No additive effect was observed in the W1282X/W1282X epithelium (Td*^CFTR^*: −5.6 ± 2.4 μA/cm^2^, Td*^CFTR^*+ETI: −5.3 ± 1.6 μA/cm^2^; **Figure 5c inset**). However, a significant increase was observed in ETI-treated Td*^CFTR^*-derived epithelium from G542X/F508del and Q493X/F508del lines (Δ: −14.8 μA/cm^2^, *p*<0.001) and F508del/F508del epithelium (Δ: −17.5 μA/cm^2^, *p*<0.05), reaching mean CFTR activity levels of −26.1 ± 2.9 and −39.1 ± 3.8 μA/cm^2^, respectively (**Figure 5c, Supplementary Table 5**).

To contextualise these gains, we analysed a cohort of non-CF participants (n=10) under matched conditions. CFTR activity in non-CF epithelium was not significantly altered by LV*^CFTR^* transduction, ETI treatment, or their combination (naive: −28.2 ± 2.0 μA/cm^2^, naive+ETI: −33.8 ± 2.8 μA/cm^2^, transduced: −31.4 ± 1.5 μA/cm^2^, Td*^CFTR^*+ETI: −34.78 ± 2.82 μA/cm^2^; **Supplementary Figure 13a-c**). When statistical comparisons were performed with a broader non-CF cohort (n=21), only the combined Td*^CFTR^*+ETI fully restored CFTR activity into the non-CF range (**Figure 5d**).

### Transplanted Rabbit Basal Cells Reconstruct Airway Epithelium and Restore Functional Mucociliary Clearance

To establish a preclinical model of airway epithelial regeneration, epithelial cells were isolated from the nasal septum of New Zealand white rabbits and expanded (in BEpiCM or Pnemacult Ex medium; **Figure 6a**). The expanded cultures formed uniform monolayers expressing the basal cell marker, p63 (**Figure 6b**). Upon air-liquid interface differentiation (*in vitro*), rabbit basal cells gave rise to functional mucociliary epithelium containing MUC5AC-positive goblet cells and acetylated tubulin-positive ciliated cells with visible beating cilia (**Figure 6c**). CFTR-mediated ion transport was confirmed by electrophysiological measurement, validating their functional capacity (**Figure 6c**). Basal cell monolayers transduced with LV^GFP^ exhibited robust GFP expression (**Figure 6d**) and were subsequently collected for transplantation into the nasal septa of a separate cohort of rabbits.

**Figure 6.**
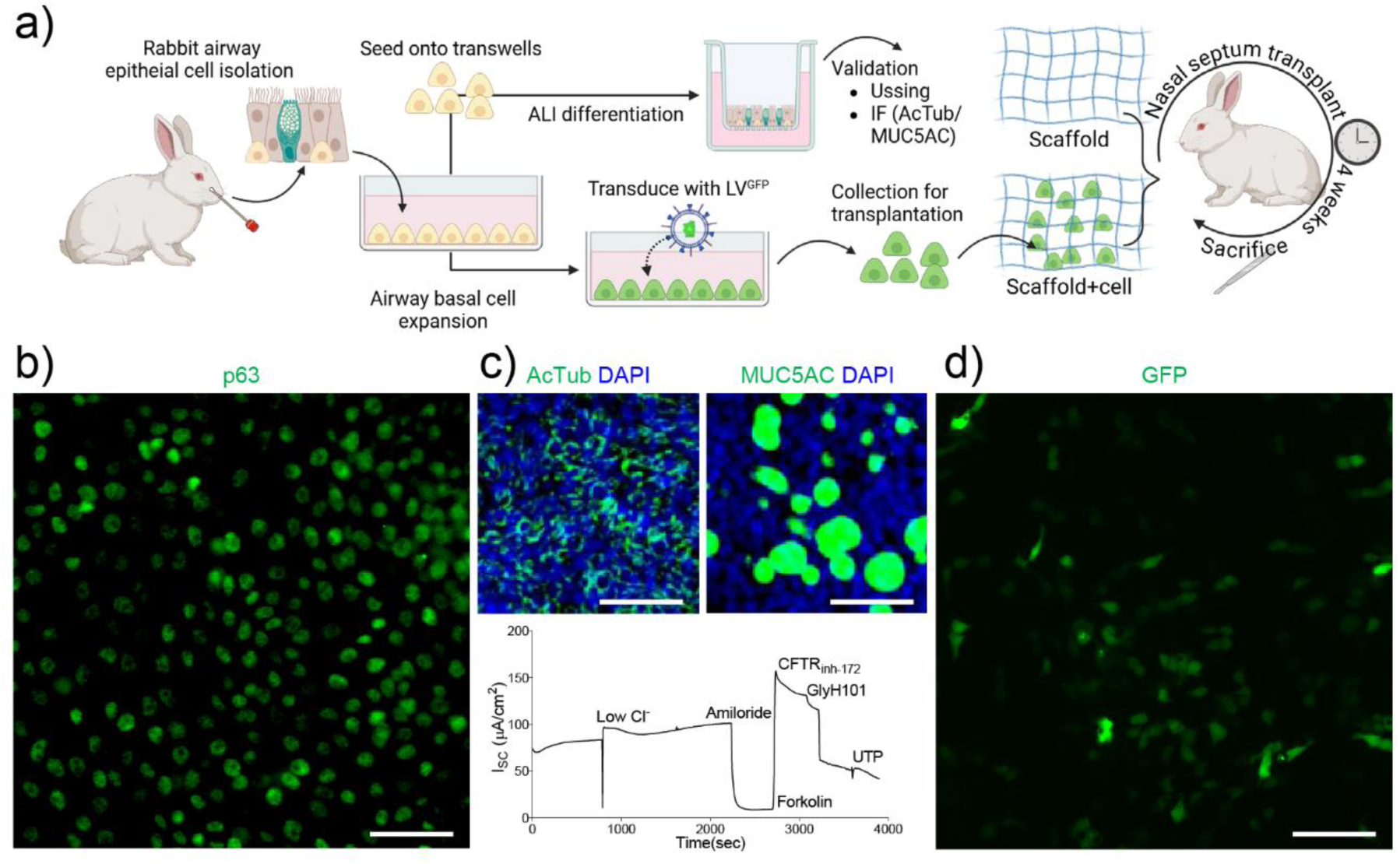
Expansion, differentiation, and LV^GFP^ transduction of rabbit basal cells for transplantation. **a)** Experimental protocol. **b)** Expansion phase: Immunofluorescence staining of basal cell marker p63 (green) at confluency. **c)** Differentiation phase: Top, (left) ciliated cells (AcTub), (right) goblet cells (MUC5AC), and bottom, CFTR-mediated short-circuit current. **d)** GFP expression following LV^GFP^ transduction. *Scale bars: 50 μm (b), 100 μm (c-d)*.

To assess engraftment *in vivo*, the nasal septum epithelium of healthy rabbits (**Figure 7a**) was completely removed by mechanical disruption, creating a receptive surface for transplantation. A cell-free tissue bioscaffold (scaffold group) or a bioscaffold seeded with LV^GFP^-transduced rabbit airway basal cells (scaffold + cell group; 5.3×10^5^/cm^2^; n= 4 per group) was then applied directly to the denuded area. This seeding density matched that used *in vitro* for ALI cultures, enabling a relevant comparison of epithelial regeneration between models.

**Figure 7.**
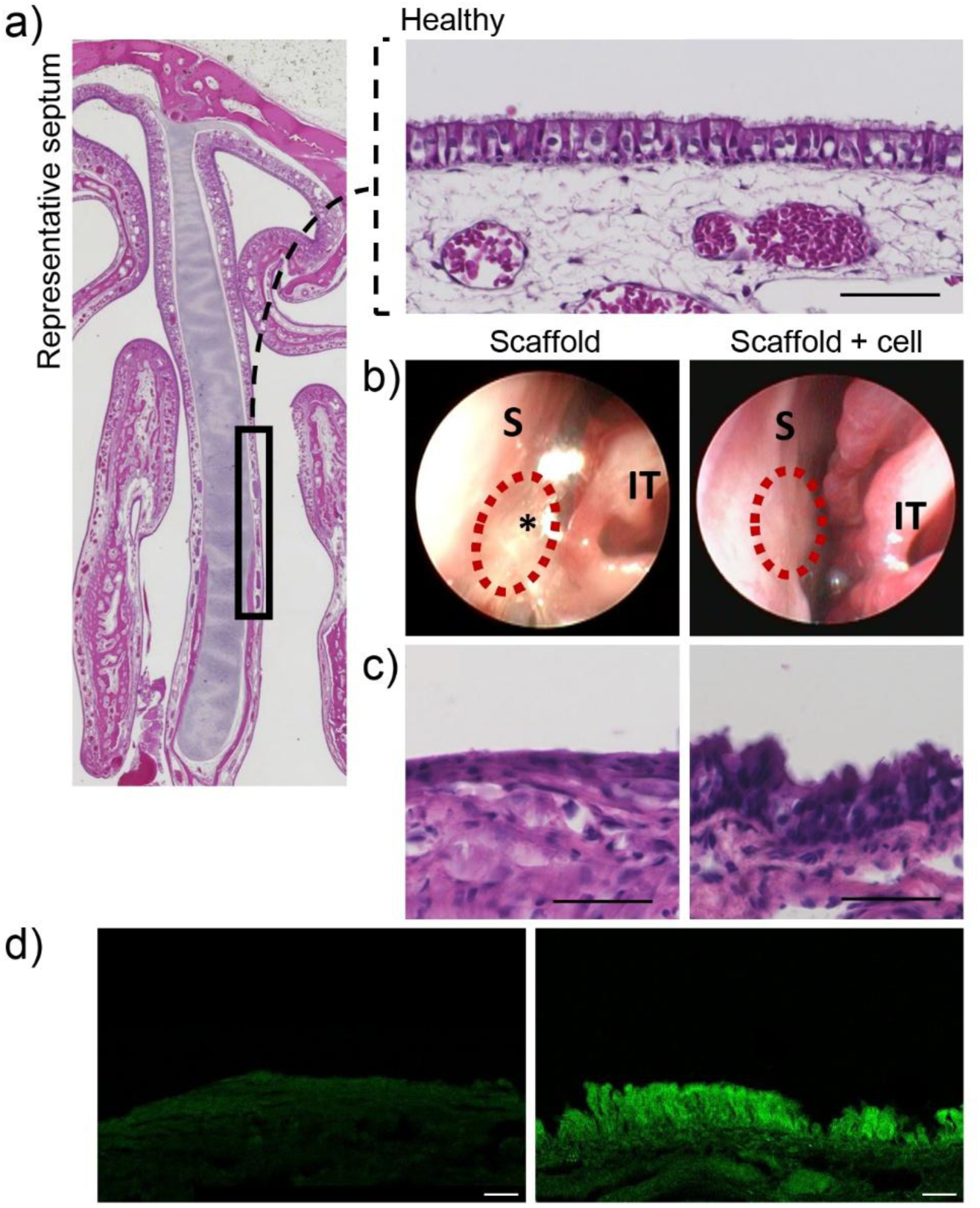
Histological assessment of epithelial regeneration in rabbit nasal septum following basal cell transplantation. **a)** Hematoxylin and eosin (H&E) staining. Diagram indicating the nasal septum region sampled (left). Representative section of the septum tissue from a healthy control rabbit (right). **b)** Representative images of the rabbit nasal septum four weeks after placement of the scaffold (left) or scaffold + cell (right). *Indicates scarring between the septum (S) and inferior turbinate (IT) complex, and the dotted circle marks the site of graft placement. **c)** Representative H&E-stained sections from rabbits receiving the scaffold (left) or scaffold + cell (right, n = 4 per group). **d)** Immunofluorescence detection of GFP signal in rabbits receiving the scaffold (left) or scaffold + cell (right). Scale bars: 50 μm.

Four weeks after transplantation, the regenerated tissue in the scaffold + cell group resembled native healthy sinus epithelium, whereas the scaffold-only group displayed epithelial thinning, structural disarray, and fibrotic scarring between the nasal septum and turbinate (**Figure 7b; Supplementary Video 1–2**). Comprehensive histological and functional analyses confirmed that rabbit airway basal cells successfully engrafted and contributed to epithelial regeneration *in vivo*. Hematoxylin and eosin (H&E) staining revealed that recipients in the scaffold + cell group developed a thicker, pseudostratified epithelium with a greater proportion of differentiated cells compared to the scaffold-only group (**Figure 7c**). Engraftment of donor cells was confirmed by detection of GFP-positive cells in the scaffold + cell group, indicating persistence and integration of LV^GFP^-transduced basal cells at the transplantation site (**Figure 7d**).

To evaluate epithelial functional following transplantation, we measured *in vivo* nasal potential difference (NPD) and performed micro-optical coherence tomography (μOCT). In the scaffold + cell group, NPD responses to CFTR agonists were significantly higher than in the scaffold-only group (– 12.6 ± 0.12 mV vs. –4.2 ± 0.6 mV; *p* < 0.001; **Figure 8a**), though both remained below values observed in healthy, uninjured controls (–33.4 ± 2.9 mV; *p* < 0.001). Importantly, because this model involves a healthy, non-CF background and no *CFTR* transgene delivery, these differences reflect variations in epithelial regeneration, not CFTR rescue. The lower NPD observed in the scaffold-only group likely results from disorganized repair, with fibrotic healing and reduced epithelial integrity, as seen histologically. In contrast, scaffold + cell grafts supported reconstitution of a more epithelial-like, pseudostratified architecture with greater numbers of differentiated cells, including CFTR-expressing ciliated and secretory lineages.

**Figure 8.**
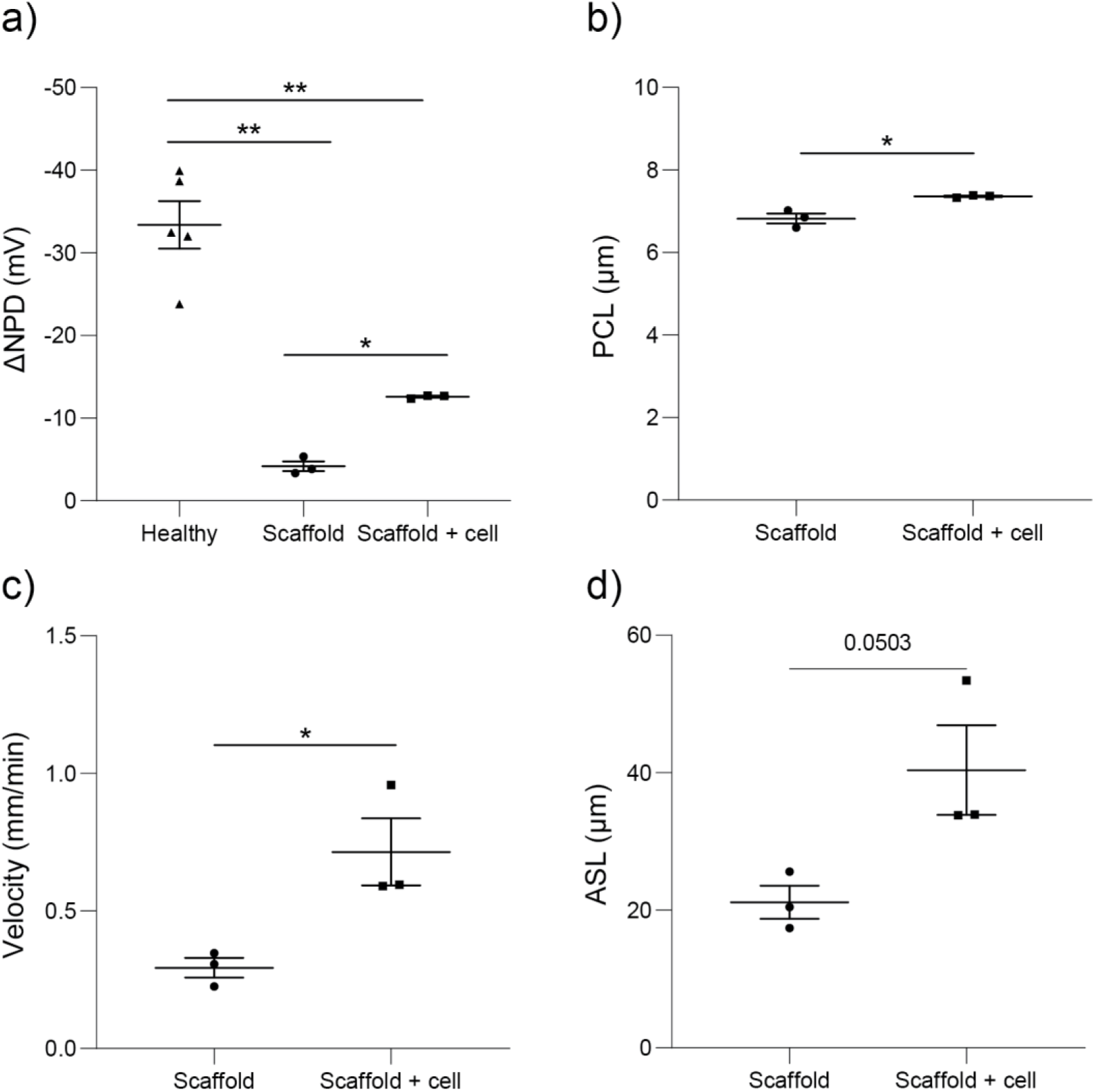
Functional restoration of rabbit airway physiology four weeks post basal cell transplantation. **a)** CFTR-dependent chloride secretion assessed by change in nasal potential difference (ΔNPD) in response to the CFTR agonist forskolin. Measurements were taken in healthy rabbits prior to mechanical disruption of the nasal septum epithelium (healthy control), and four weeks after transplantation with either scaffold-only or scaffold+cell treatment. **b)** Periciliary liquid layer (PCL) depth, **c)** mucociliary transport (MCT) velocity and **d)** airway surface liquid (ASL) depth, all measured by micro-optical coherence tomography (µOCT). Data are shown as mean ± SEM (n=3 per group). Statistical comparisons were performed using unpaired t-tests. \**p*<0.05, \*\**p*<0.01.

μOCT imaging further supported these findings. The scaffold + cell group exhibited significantly improved mucociliary parameters, including increased periciliary liquid depth (PCL: 7.36 ± 0.02 μm vs. 6.82 ± 0.12 μm; *p* < 0.01; **Figure 8b**) and faster mucociliary transport (MCT: 0.72 ± 0.12 mm/min vs. 0.29 ± 0.04 mm/min; *p* = 0.05; **Figure 8c**). Airway surface liquid depth also trended higher (ASL: 40.4 ± 6.5 μm vs. 21.2 ± 2.4 μm; *p* = 0.0503; **Figure 8d**). These outcomes are consistent with improved epithelial regeneration and functional recovery of mucociliary clearance driven by the transplanted basal cells.

Collectively, these results suggest that transplanted basal cells contribute to restoration of native epithelial function after injury, likely by accelerating re-epithelialization and maintaining cell types essential for airway physiology, including CFTR-expressing cells.

## Discussion

We demonstrate that LV *CFTR* gene addition restores CFTR function in differentiated human airway basal cells *in vitro*. Corrected cells formed well-structured pseudostratified epithelium with improved CFTR activity across multiple CF genotypes. In parallel, we used LV^GFP^-labelled, non-CF, donor-derived rabbit basal cells to assess epithelial engraftment and mucociliary regeneration *in vivo*. As no *CFTR* transgene was delivered, these *in vivo* findings reflect epithelial repair capacity following injury, rather than CFTR correction or disease modelling. Together, our results establish a two-part platform: functional CFTR restoration in human cells *in vitro*, and airway regeneration from basal cells in a relevant *in vivo* model.

Building on foundational work in sinonasal [14], tracheal [13], and bronchial [12–14, 16] epithelial models, we show that LV mediated CFTR delivery does not compromise epithelial differentiation, structural integrity, or cellular composition. Transduced cells differentiated into well-organized pseudostratified epithelium with preserved tight junctions, normal ciliary function, and global proteome stability across both intracellular and secreted compartments, which underscores the minimal off-target impact of CFTR transgene expression. Notably, CFTR activity in Td*^CFTR^*-derived epithelium matched or exceeded that of ETI-treated naive cells, and additive effects were observed when both therapies were combined—reaching non-CF levels in modulator-eligible participants.

The ability to restore CFTR activity even in participants with nonsense mutations, such as W1282X/W1282X, is especially significant. Approximately 10% of pwCF remain without effective modulators [3]; gene addition offers a durable, mutation-agnostic approach to overcome this gap. Moreover, the observed additive effect between gene addition and ETI in ETI-responsive participants suggests these approaches are not mutually exclusive but may be synergistic—offering a path toward fully normalized CFTR function, even in modulator-responsive patients.

Transduction efficiency varied widely (1.2–40.2%) but strongly correlated with CFTR activity (ρ = - 0.92), emphasizing the importance of achieving threshold-level gene delivery. Our observation that even modest transduction levels conferred benefit, aligns with clinical data suggesting that restoring CFTR function to as little as 10–25% of normal CFTR function can significantly ameliorate disease severity [30]. Strategies to enhance LV uptake, such as G2-phase modulation [31], use of LentiBOOST [32], or inhibition of autophagy/proteolysis [33, 34], warrant systematic evaluation in airway basal cells. However, increasing the MOI must be carefully balanced to preserve cell viability and avoid integration-related toxicity. This is especially important since increasing CFTR mRNA does not necessarily yield improved CFTR-mediated Cl⁻ transport [35]. Defining the therapeutic window for *CFTR* addition remains essential for clinical translation.

Unlike transient mRNA or AAV-based approaches, LV vectors stably integrate and support long-term expression. Our use of the EF1α promoter, previously shown to outperform PGK in driving CFTR activity [12], enabled broad epithelial transgene expression, including in ciliated and secretory lineages. This heterogeneity reflects basal cell plasticity and supports the capacity of transduced cells to functionally regenerate the epithelium. While long-term persistence of CFTR activity under inflammatory or stress conditions remains to be tested, our proteomic and functional data show no loss of epithelial identity or CFTR function up to the endpoints evaluated. Intriguingly, we observed a correlation between CFTR restoration and elevated ciliary beat frequency, suggesting that gene correction may also improve mucociliary clearance—particularly relevant in advanced lung disease, where impaired mucus transport contributes to chronic infection and inflammation.

While gene editing offers the potential for precise correction, current approaches face challenges related to editing efficiency, cell fitness, and delivery scalability [14–16]. Gene addition, in contrast, offers a modular and mutation-agnostic platform with clinical scalability, well suited for both autologous and allogeneic cell therapy strategies. In our study, CFTR restoration levels varied across donors despite uniform vector design, suggesting that differences in cellular health, epigenetic landscape, or integration site may influence transduction efficiency outcomes. Comparisons across studies are further complicated by differences in vector design and promoter choice, which can significantly impact CFTR expression and Cl⁻ transport [35]. Moving forward, systematically dissecting the contribution of these variables will be essential for standardizing gene addition strategies and ensuring consistent therapeutic efficacy across diverse patient populations.

While the *in vitro* arm of our study directly demonstrated functional CFTR rescue, the *in vivo* model served to assess the capacity of airway basal cells to regenerate epithelium and restore mucociliary architecture in a non-diseased setting. A key translational advance is our demonstration of functional engraftment in the rabbit sinus model. Several groups have made significant progress toward airway cell therapies, including studies showing engraftment of airway basal cells in the trachea of mouse models [17, 18, 36]. However, most previous efforts have focused on histological or basic functional readouts, such as CBF or oxygen saturation, with limited comprehensive assessment of mucociliary function *in vivo*.

The paranasal sinuses provide a clinically actionable site for cell therapy delivery due to their anatomical similarity to the lower airway epithelium, surgical accessibility, and central role in perpetuating infection, including in lung transplant recipients, where sinus-derived pathogens can recolonize allografts [23]. Our model may also be valuable beyond CF, offering a platform to evaluate regenerative therapies for other upper airway disorders, including chronic rhinosinusitis, which affects over 10% of the global population and lacks durable treatments for epithelial dysfunction. Additionally, sinus-directed repair strategies could benefit patients undergoing endoscopic sinus surgery, where epithelial regeneration is critical for post-operative recovery and long-term symptom resolution. In our study, transplanted rabbit basal cells regenerated a pseudostratified mucociliary epithelium with functional evidence of engraftment demonstrated by NPD, PCL depth, and MCT velocity.

While encouraging, these *in vivo* studies represent an early proof-of-concept. Long-term persistence, immunogenicity, and scalability remain open challenges. This may be resolved by autologous cell transplant or use of a universal donor. Prior work has shown that suppressing TGF-β signalling with pirfenidone enhances engraftment in murine models [18]. Future strategies could also include transient immunosuppression, co-transplantation with niche-stabilizing cells, or scaffold-based delivery systems tailored for immune modulation. Here, we empirically selected a cell dose that recapitulated ALI culture density; however, dose-ranging studies and Good Manufacturing Practice-compliant scale-up protocols will be critical to enable clinical translation.

Our findings support a flexible therapeutic model: as a standalone treatment for individuals ineligible for modulators, as an adjunct therapy to optimize outcomes for those on ETI, and as a targeted sinus-directed intervention for lung transplant recipients no longer eligible for modulator therapy but vulnerable to upper airway-driven reinfection. This modularity positions *CFTR* gene and cell therapy as a versatile addition to the treatment landscape.

## Acknowledgments

We thank the study participants and their families for their invaluable contributions. We appreciate the assistance from Sydney Children’s Hospitals Randwick respiratory team for biospecimen collection. We thank Nihan Turgutoglu for assistance with primary cell culture. Technical assistance was provided by the Bioanalytical Mass Spectrometry Facility, Katharina Gaus Light Microscopy Facility and Dr Emma Johansson Beves at the Flow Cytometry Facility of Mark Wainwright Analytical Centre at UNSW Sydney, Jason Gummow of the Functional Genomics South Australia Core Facility (FGSA), University of Adelaide, and Dr Jeffery McArthur from the Victor Chang Cardiac Researcher Institute Innovation Centre. Advice regarding the statistical analysis was provided by Nancy Briggs from Stats Central, UNSW. Figures 1 and 6a were created in Biorender (Waters, S. (2025) https://BioRender.com/qmr4e9h, https://BioRender.com/avoqxzn).

## References

1. Rommens JM. Cystic Fibrosis Mutation Database 2011. http://www.genet.sickkids.on.ca/Home.html. Date last updated: April 25 2011. Date last accessed: June 27 2025.

2. Grasemann H, Ratjen F. Cystic Fibrosis. N Engl J Med. 2023;389(18):1693–707.

3. Middleton PG, Mall MA, Drevinek P, et al. Elexacaftor-Tezacaftor-Ivacaftor for Cystic Fibrosis with a Single Phe508del Allele. N Engl J Med. 2019;381(19):1809–19.

4. Allan KM, Farrow N, Donnelley M, et al. Treatment of Cystic Fibrosis: From Gene-to Cell-Based Therapies. Front Pharmacol. 2021;12:639475.

5. Rock JR, Onaitis MW, Rawlins EL, et al. Basal cells as stem cells of the mouse trachea and human airway epithelium. Proc Natl Acad Sci U S A. 2009;106(31):12771–5.

6. Montoro DT, Haber AL, Biton M, et al. A revised airway epithelial hierarchy includes CFTR-expressing ionocytes. Nature. 2018;560(7718):319-24.

7. Davies JC, Polineni D, Boyd AC, et al. Lentiviral Gene Therapy for Cystic Fibrosis: A Promising Approach and First-in-Human Trial. Am J Respir Crit Care Med. 2024;210(12):1398–408.

8. Alton EW, Beekman JM, Boyd AC, et al. Preparation for a first-in-man lentivirus trial in patients with cystic fibrosis. Thorax. 2017;72(2):137–47.

9. McCarron A, Farrow N, Cmielewski P, et al. Breaching the Delivery Barrier: Chemical and Physical Airway Epithelium Disruption Strategies for Enhancing Lentiviral-Mediated Gene Therapy. Front Pharmacol. 2021;12:669635.

10. Farrow N, Donnelley M, Cmielewski P, et al. Role of Basal Cells in Producing Persistent Lentivirus-Mediated Airway Gene Expression. Hum Gene Ther. 2018;29(6):653–62.

11. Carpentieri C, Farrow N, Cmielewski P, et al. The Effects of Conditioning and Lentiviral Vector Pseudotype on Short- and Long-Term Airway Reporter Gene Expression in Mice. Hum Gene Ther. 2021;32(15-16):817–27.

12. Marquez Loza LI, Cooney AL, Dong Q, et al. Increased CFTR expression and function from an optimized lentiviral vector for cystic fibrosis gene therapy. Mol Ther Methods Clin Dev. 2021;21:94–106.

13. Cooney AL, Thurman AL, McCray PB, Jr., et al. Lentiviral vectors transduce lung stem cells without disrupting plasticity. Mol Ther Nucleic Acids. 2021;25:293–301.

14. Vaidyanathan S, Salahudeen AA, Sellers ZM, et al. High-Efficiency, Selection-free Gene Repair in Airway Stem Cells from Cystic Fibrosis Patients Rescues CFTR Function in Differentiated Epithelia. Cell Stem Cell. 2020;26(2):161–71 e4.

15. Vaidyanathan S, Baik R, Chen L, et al. Targeted replacement of full-length CFTR in human airway stem cells by CRISPR-Cas9 for pan-mutation correction in the endogenous locus. Mol Ther. 2022;30(1):223–37.

16. Suzuki S, Crane AM, Anirudhan V, et al. Highly Efficient Gene Editing of Cystic Fibrosis Patient-Derived Airway Basal Cells Results in Functional CFTR Correction. Mol Ther. 2020;28(7):1684–95.

17. Ma L, Thapa BR, Le Suer JA, et al. Airway stem cell reconstitution by the transplantation of primary or pluripotent stem cell-derived basal cells. Cell Stem Cell. 2023;30(9):1199–216 e7.

18. Ma Q, Ma Y, Dai X, et al. Regeneration of functional alveoli by adult human SOX9(+) airway basal cell transplantation. Protein Cell. 2018;9(3):267–82.

19. Sun F, Cheng L, Guo H, et al. Application of autologous SOX9(+) airway basal cells in patients with bronchiectasis. Clin Respir J. 2020;14(9):839–48.

20. Wang Y, Meng Z, Liu M, et al. Autologous transplantation of P63(+) lung progenitor cells for chronic obstructive pulmonary disease therapy. Sci Transl Med. 2024;16(734):eadi3360.

21. Yan J, Zhang W, Feng Y, et al. Autologous transplantation of P63(+) lung progenitor cells in patients with bronchiectasis: A randomized, single-blind, controlled trial. Cell Rep Med. 2024;5(11):101819.

22. Liu Z, Zheng Q, Li Z, et al. Epithelial stem cells from human small bronchi offer a potential for therapy of idiopathic pulmonary fibrosis. EBioMedicine. 2025;112:105538.

23. Choi KJ, Cheng TZ, Honeybrook AL, et al. Correlation between sinus and lung cultures in lung transplant patients with cystic fibrosis. Int Forum Allergy Rhinol. 2018;8(3):389–93.

24. Schogler A, Blank F, Brugger M, et al. Characterization of pediatric cystic fibrosis airway epithelial cell cultures at the air-liquid interface obtained by non-invasive nasal cytology brush sampling. Respir Res. 2017;18(1):215.

25. Ozcan KM, Ozcan I, Selcuk A, et al. Comparison of Histopathological and CT Findings in Experimental Rabbit Sinusitis. Indian J Otolaryngol Head Neck Surg. 2011;63(1):56–9.

26. Allan KM, Wong SL, Fawcett LK, et al. Collection, Expansion, and Differentiation of Primary Human Nasal Epithelial Cell Models for Quantification of Cilia Beat Frequency. J Vis Exp. 2021(177):e63090.

27. Wong SL, Kardia E, Vijayan A, et al. Molecular and Functional Characteristics of Airway Epithelium under Chronic Hypoxia. Int J Mol Sci. 2023;24(7).

28. Cho DY, Zhang S, Skinner DF, et al. Ivacaftor restores delayed mucociliary transport caused by Pseudomonas aeruginosa-induced acquired cystic fibrosis transmembrane conductance regulator dysfunction in rabbit nasal epithelia. Int Forum Allergy Rhinol. 2022;12(5):690–8.

29. Ricana CL, Lyddon TD, Dick RA, et al. Primate lentiviruses require Inositol hexakisphosphate (IP6) or inositol pentakisphosphate (IP5) for the production of viral particles. PLoS Pathog. 2020;16(8):e1008646.

30. Ramalho AS, Beck S, Meyer M, et al. Five percent of normal cystic fibrosis transmembrane conductance regulator mRNA ameliorates the severity of pulmonary disease in cystic fibrosis. Am J Respir Cell Mol Biol. 2002;27(5):619–27.

31. Zhang S, Joseph G, Pollok K, et al. G2 cell cycle arrest and cyclophilin A in lentiviral gene transfer. Mol Ther. 2006;14(4):546–54.

32. Lo Presti V, Cornel AM, Plantinga M, et al. Efficient lentiviral transduction method to gene modify cord blood CD8(+) T cells for cancer therapy applications. Mol Ther Methods Clin Dev. 2021;21:357–68.

33. Santoni de Sio FR, Cascio P, Zingale A, et al. Proteasome activity restricts lentiviral gene transfer into hematopoietic stem cells and is down-regulated by cytokines that enhance transduction. Blood. 2006;107(11):4257–65.

34. Wang CX, Sather BD, Wang X, et al. Rapamycin relieves lentiviral vector transduction resistance in human and mouse hematopoietic stem cells. Blood. 2014;124(6):913–23.

35. Farmen SL, Karp PH, Ng P, et al. Gene transfer of CFTR to airway epithelia: low levels of expression are sufficient to correct Cl-transport and overexpression can generate basolateral CFTR. Am J Physiol Lung Cell Mol Physiol. 2005;289(6):L1123–30.

36. Hawkins FJ, Suzuki S, Beermann ML, et al. Derivation of Airway Basal Stem Cells from Human Pluripotent Stem Cells. Cell Stem Cell. 2021;28(1):79–95 e8.

