## Supplemental Material for "Gene-Corrected Basal Cells Restore CFTR In Vitro; Transplants Regenerate Epithelium in a Preclinical Sinus Model"

### Supplementary Material

### Material and Methods

*Vector production*

VSV-G–pseudotyped HIV-1–based LV vectors expressing V5-tagged *CFTR* ($LV^{CFTR}$) or enhanced green fluorescent protein ($LV^{eGFP}$) were driven by the EF1α promoter and produced and titered by the Gene Silencing and Expression Facility (Robinson Research Institute, The University of Adelaide, Adelaide, Australia). $LV^{CFTR}$ titres were quantified by qPCR ($7.45\times10^7$ – $3.45\times10^8$ IU/mL), and $LV^{eGFP}$ titres by flow cytometry ($1.66\times10^9$ IU/mL).

*Participant airway epithelial cell collection and expansion*

Participants were recruited under the CF AVATAR study (HREC/16/SCHN/120) with written guardian consent. Human airway epithelial cells (hAECs) were collected by nasal inferior turbinate brushing (McFarlane Medical, Ringwood, Australia) or from bronchoalveolar lavage fluid as previously described [1]. Samples included five participants with a Class I *CFTR* mutation (G542X or Q493X, one with homozygous W1282X) and seven homozygous for the F508del *CFTR* mutation (three also with matched bronchial epithelial cells (hBECs); **Supplementary Table 1**). All CF participants were pancreatic insufficient (ages 1.7-14.9 years). Control nasal epithelial cells were also collected from ten non-CF participants (ages 0.9-16.2 years).

hAECs were expanded as described previously [2]. Briefly, cells were seeded (5,000 cells/$cm^2$) onto collagen I-coated (Advanced Biomatrix, Carlsbad, CA) flasks pre-seeded with irradiated NIH/3T3 feeder cells [2, 3]. Expansion was performed using conditionally reprogramming cell media with ROCK Inhibitor Y-27632 (10 μM; Selleckchem, Houston, USA, S1049) to generate airway basal cell monolayers. At confluence, monolayers were dissociated via differential concentration trypsinisation and cryopreserved.

*Transduction of airway basal cells*

Passage one airway basal cells were seeded (52,000 cells/$cm^2$) onto collagen I-coated 6-well plates pre-seeded with irradiated NIH/3T3 feeder cells. At 70% confluence, cells (~$3.0\times10^6$) were transduced with $LV^{CFTR}$ or $LV^{eGFP}$ at a MOI of 10 or 20. Untransduced (naive) controls were cultured in parallel. After overnight incubation at 37°C and 5% $CO_2$, cells were collected and used for downstream experiments and cryopreserved. For repeat transduction experiments, cryopreserved $LV^{CFTR}$-transduced cells were re-seeded (52,000 cells/$cm^2$) onto collagen I-coated 6-well plates pre-seeded with irradiated NIH/3T3 feeder cells and at 70% confluence, and a second $LV^{CFTR}$ transduction (MOI 10) was performed. Control cells were cultured in parallel without re-transduction.

*Immunofluorescence labelling, imaging and quantitative data processing of basal cell monolayers*

Immunolabelling was automated using OT-2 robotics (Opentrons, Brooklyn, USA). Cells were fixed in 4% paraformaldehyde (20 min), permeabilised with 1% Triton X-100 (10 min, room temperature), and blocked with Intercept Buffer (LI-COR, Lincoln, USA, LCR-927-70001). Samples were incubated with anti-V5-Tag (1:500; D3H8Q, Cell Signalling Technology, Danvers, USA, 13202) for 2 h, followed by Alexa Fluor 555–conjugated secondary antibody (1:1000; Cell Signalling Technology, 4413) for 30 min. DAPI (1:2000; Sigma-Aldrich, Saint Louis, USA, D9542) and Phalloidin-Atto 647N (1:2000; ATTO-TEC, Siegen, Germany, AD647N) were used to stain nuclei and actin.

Confocal images were acquired on a Nikon AX-R (Nikon, Tokyo, Japan) using a 20× Plan Apochromat air objective (NA 0.75), 2× zoom, and resonance scanning (2048×2048 pixels). A 4-channel sequential detection mode was used: 405/647 nm (pass 1), 488 nm (pass 2), and 555 nm (pass 3). Images were saved as 12-bit TIFFs for analysis.

Cell segmentation and feature extraction were performed using CellProfiler v4.2.1 (Broad Institute, MIT, Cambridge, USA) [4] with the minimum cross-entropy algorithm, propagating from DAPI-stained nuclei to Phalloidin-defined cell borders. 574 features per cell were extracted, covering morphology and antibody signal metrics. Data were analysed in Konstanz Information Miner software v4.7.4 (KNIME, Zurich, Switzerland) with z-score normalisation, followed by principal component analysis (PCA, 97% variance retention) and t-distributed stochastic neighbour embedding (t-SNE) visualisation [5].

*Flow cytometry*

$LV^{CFTR}$ transduction efficiency was assessed by V5 expression. Cells were fixed in 4% paraformaldehyde, permeabilized with 90% methanol, stained with anti-V5-Tag (D3H8Q, Cell Signalling Technology, Danvers, USA, 13202) for 1 h, then with Alexa Fluor 488-conjugated goat anti-rabbit secondary antibody (Thermo Fisher Scientific, Waltham, USA, A11034) for 30 min. A total of $0.25\text{-}0.5\times10^6$ cells were resuspended in FACS buffer (20% FBS and 2mM EDTA in PBS) and analysed on a BD FACSAria II flow cytometer (BD Biosciences, Franklin Lakes, USA). At least 50,000 events were recorded per sample. Unstained and secondary-only cells were used as negative control. Transduction efficiency was calculated as ([$V5^+$ cells/total cells] × 100). Data were analysed using BD FACSDiva software (BD Biosciences, Franklin Lakes, USA).

*RT-qPCR*

Samples were prepared as previously described [6]. RNA was extracted using TRIzol (Thermo Fisher Scientific, Waltham, USA) and purified with the RNeasy Mini Kit (QIAGEN, Hilden, Germany). RNA concentration was measured, and 120 ng was used for cDNA synthesis (M-MLV Reverse Transcriptase and random hexamer priming). The exogenous *CFTR* transgene and endogenous F508del *CFTR* were detected using primer pairs from Clarke et al. (2019) [7]. C(t) values were normalized to ACTB (F: AGAAAATCTGGCACCACACC; R: AGAGGCGTACAGGGATAGCA), and expression was calculated using the $2^{(-\Delta\Delta C_t)}$ method [6, 7], with F508del as an internal reference.

*Whole cell patch-clamp electrophysiology*

$LV^{CFTR}$-transduced cells were cultured for two weeks, then dissociated for automated patch clamp on the SyncroPatch 384PE (Nanion Technologies, Munich, Germany). A ramp protocol (-100 to +100 mV; 10 mV/ms) was applied every 10 s. CFTR activation used forskolin (10 µM; Sigma-Aldrich, Saint Louis, USA, F6886) and VX-770 (10 µM; Selleckchem, Houston, USA, S1144); inhibition was induced with Inh-172 (10 µM; Sigma-Aldrich, Saint Louis, USA, C2992). Current density was normalised to capacitance (pA/pF). The percentage of CFTR-active cells was calculated as (cells with Inh-172 current/total with proper cell catch).

*Mucociliary differentiation of airway basal cells at air liquid interface (ALI)*

Naive and transduced basal cells (250,000 cells/ membrane) were seeded onto collagen I-coated 6.5 mm 0.4 µm Transwell polyester membranes (Sigma-Aldrich, Saint Louis, USA, CLS3470), and cultured as previously described [2, 6]. In brief, cells were expanded in PneumaCult Ex Plus Expansion media (STEMCELL Technologies, Vancouver, Canada, 05040) until confluence, then differentiated in PneumaCult ALI media (STEMCELL Technologies, Vancouver, Canada, 05001) for 21-28 days until a mature pseudostratified epithelium with beating cilia was established. Media was changed every second day; mucus was removed weekly by PBS washing.

*Whole mount immunofluorescence*

ALI cultures derived from four of ten CF participants were fixed (4% paraformaldehyde, 30 min, RT), permeabilised (0.5% Triton X-100) and blocked in IF buffer, followed by primary antibody incubation for V5 tag, acetylated tubulin, MUC5AC, and p63 (**Supplementary Table 2**). Alexa Fluor-conjugated secondary antibodies, phalloidin and DAPI were used for counterstaining. Confocal images were acquired on a Leica TCS SP8 DLS confocal microscope (63×/1.4 objective, Leica Microsystems, Wetzlar, Germany) and processed in ImageJ (National Institutes of Health, Bethesda, USA), adjusting the dynamic range of intensity for target visualisation.

*Sample preparation for Mass Spectrometry*

Apical secretome was collected from ALI cultures (n=3 F508del/F508del) with PBS washes, precipitated in cold 30% TCA/acetone, centrifuged (12000 × g), then reconstituted in 2% SDC/50 mM

Tris-HCl buffer. Lysates from naive and transduced cells were processed via RIPA lysis and sonication, prior to protein concentration determination using a Pierce BCA Protein Assay Kit (Thermo Fisher Scientific, Waltham, USA). Samples were reduced, alkylated, digested with trypsin, and purified by strong cation exchange, then reconstituted in formic acid/acetonitrile mix as previously described [1].

*Mass Spectrometry*

Proteolytic peptide samples were separated by nanoLC using an Ultimate nanoRSLC UPLC system with an autosampler (Dionex, Amsterdam, Netherlands). Peptides were eluted at 200 nL/min using a linear gradient from $H_2O$:$CH_3CN$ (98:2, 0.1% formic acid) to $H_2O$:$CH_3CN$ (64:36, 0.1% formic acid) over 60 min (apical secretome) or 90 min (cell lysate). Eluted peptides were ionized in positive ion mode nano-ESI (2000 V) using a low-volume titanium union with the tip positioned ~0.5 cm from the heated capillary (275°C) of a Tribrid Fusion Lumos mass spectrometer (Thermo Fisher Scientific, Waltham, USA). The mass spectrometry proteomics data have been deposited in the ProteomeXchange Consortium via the PRIDE [8] partner repository (Dataset ID: PXD034191), available for review via: username:, password: dXIVwj4H.

*Protein identification, quantification, and statistical analysis*

LC-MS/MS raw files were analysed using MaxQuant (v1.6.2.10.43) with sequence searches performed using Andromeda [9]. Label-free quantification was performed using the MaxLFQ algorithm [10]. Search parameters included: carbamidomethyl (C) as a fixed modification; oxidation (M) and N-terminal protein acetylation as variable modifications; and enzyme specificity was trypsin with up to two missed cleavages. Peaks were searched against the human Swiss-Prot database (July 2021 release, 20588 sequences) with a minimum peptide length of 7. MaxLFQ analyses were performed with default parameters, enabling “fast LFQ”. Protein and peptide false discovery rates (FDR) were set at 1%, and only non-contaminant proteins identified from ≥2 unique peptides were used for downstream analysis.

Differential protein expression analysis was conducted using the R package DEP (v1.10.0) [11] within R using RStudio. Proteins detected in ≥ 50% of samples were retained, and missing values were imputed using BPCA. Proteins were considered differentially abundant if they had a fold change > 1.2 and *p*-value < 0.05. Functional analysis of differentially abundant proteins was performed using Ingenuity Pathway Analysis (IPA; QIAGEN, Hilden, Germany) [12]. Proteomic cell-type marker analysis was performed as described previously [3].

*Cilia beating frequency (CBF) measurements*

Cilia beating was recorded in ALI cultures at 21-25 days as described previously [2]. Imaging was conducted in an environmental chamber (37°C, 5% $CO_2$, 85% relative humidity) following a 30-min equilibration period. Fast frame rate imaging (>100 Hz) was performed using an Andor Zyla 4.2

sCMOS camera (Oxford Instruments, Abingdon, UK) connected to a Nikon Eclipse Ti2-E live-cell imaging microscope (Nikon, Tokyo, Japan) with a 20×/0.8 long working distance objective. Fast time-lapse images (1000 frames; ~333 frames per second) were acquired from randomly sampled 512 × 512 pixels regions across triplicate filters. CBF was analysed using a previously developed custom script [2].

*CFTR modulator pre-treatment*

Naive and transduced ALI cultures were incubated basolaterally with elexacaftor (VX-445, 3 μM; Selleckchem, Houston, USA, S8851) and tezacaftor (VX-661, 18 μM; Selleckchem, Houston, USA, S7059) for 48 h prior to Ussing chamber assays. Following pre-treatment, ivacaftor (VX-770, 10 μM; Selleckchem, Houston, USA, S1144) was added during the Ussing protocol to complete the ETI treatment.

*Short circuit current measurements in Ussing chambers*

Naive and transduced ALI cultures were mounted in circulating Ussing chambers (VCC MC8; Physiologic Instruments, San Diego, USA). Three independent ALI cultures were tested per participant per condition. Short-circuit current ($I_{SC}$, μA/cm$^2$) was measured under asymmetric $Cl^-$ Ringer's buffer (bicarbonate free) as described previously [3]. After recording baseline current for 30 min, cultures were sequentially treated with: 100 μM apical amiloride (Sigma-Aldrich, Saint Louis, USA, A7410), DMSO (naive) or 10 μM apical–VX-770 (ETI-treated), 10 μM basolateral forskolin, 30 μM apical $CFTR_{inh}$-172 and 100 μM apical ATP (Sigma-Aldrich, Saint Louis, USA, A2383). Data were recorded using Acquire and Analysis 2.3 software (Physiologic Instruments, San Diego, USA). Changes in $I_{SC}$ following forskolin and $CFTR_{inh}$-172 indicated CFTR activity. Filter and solution resistance (without cells) were subtracted from all measurements. CF ALI cultures were compared to non-CF CFTR activity levels in ALI cultures tested in this study (n=10) and in our previous work (n=11) [13].

*In vitro rabbit airway epithelial cell culture, validation and transduction*

This study was approved by the Institutional Animal Care and Use Committee (IACUC) at the University of Alabama at Birmingham (UAB) under protocol IACUC-09584. Rabbit airway epithelial cells were collected from the nasal septum immediately after euthanasia and dissociated as described previously [14]. Cells were expanded in Pneumacult-Ex Medium (STEMCELL Technologies, Vancouver, Canada, 05008) or Bronchial Epithelial Cell Medium (BEpiCM, ScienCell, Carlsbad, USA, 3211) to establish basal cell monolayers [3]. For validation of differentiation capacity, cells were then dissociated, seeded onto Transwell filters (Corning, Carlsbad, USA, 3470) at $2\times10^5$ cells/filter, and differentiated under ALI conditions in ALI medium until maturity. Fully differentiated ALI cultures were mounted in Ussing chambers and short-circuit current was measured as described previously [14].

For transduction and tagging of cells, passage one rabbit airway basal cells were seeded into coated 24-well plates (0.5-1×$10^5$ cells/well). At 75-80% confluence, cells were transduced with 1.8×$10^4$ TU/ml of MISSION pLKO.1-puro-CMV-TurboGFP Positive Control Transduction Particles (Sigma-Aldrich, Saint Louis, USA, SHC003VN).

*In vivo rabbit model engraftment and transplant*

This study was approved by the Institutional Animal Care and Use Committee (IACUC) at the University of Alabama at Birmingham (UAB) under protocol IACUC-22329. Female New Zealand White rabbits (3–4 kg), confirmed Pasteurella-free, were used. Rabbits were acclimated to the animal facility for at least one week. Anaesthesia was administered in a warm environment to ensure comfort, using a combination of ketamine (20 mg/kg; MWI, Boise, ID), dexdomitor (0.25 mg/kg; Zoetis Inc., Parsippany, USA), buprenorphine (0.03 mg/kg; Reckitt Benckiser Pharmaceuticals Inc., Parsippany, USA), and carprofen (5 mg/kg; Zoetis Inc., Parsippany, USA). To confirm the absence of pre-existing infections or abnormal lesions, rabbits underwent nasal endoscopic examination using a 1.7 mm, 30-degree scope (Karl Storz, Tuttlingen, Germany).

A 25 µl suspension of GFP-transduced rabbit cells (1.6×$10^5$) was seeded onto a 6 mm diameter Myriad Matrix Soft Tissue Bioscaffold (Aroa Biosurgery Limited, San Diego, USA, 09421906017649), clinically used for plastic and reconstructive surgery and wound repair [15, 16]. The cell-seeded scaffold was placed on the exposed rabbit nasal septal cartilage (n=4), while the control rabbits received a scaffold with no cells (n=4). Post transplantation assessments included *ex vivo* micro-optical coherence tomography (µOCT) imaging (n=1 per group) and NPD measurements, followed by humane kill at four weeks for histological staining (n=3 per group).

*Histological staining*

Sinus tissues were collected, fixed in 4% paraformaldehyde, and subjected to EDTA decalcification before flash freezing. Samples were stored at -80°C until further processing. GFP expression was assessed in unstained tissue sections using an A1R-HD25 Confocal Microscope (Nikon, Tokyo, Japan). Hematoxylin and eosin (H&E) staining was performed on parallel sections (n=3 per group), with images captured using a Keyence BZ-X800 microscope (Keyence, Osaka, Japan).

*Immunofluorescence staining*

Monolayer cultures and fully differentiated ALI cultures (day 15) were fixed in 4% paraformaldehyde, then neutralized, permeabilized, and blocked before primary antibody incubation: P63 (Santa Cruz, Dallas, USA, sc-25268) for 24 h, MUC5AC (Abcam, Cambridge, UK, ab3649) for 16 h, and acetylated tubulin (Abcam, Cambridge, UK, ab56676) for 24 h at 4°C. Following incubation, samples were washed with IF buffer, incubated with Alexa Fluor-conjugated secondary antibodies (Invitrogen,

Waltham, USA, A-11029) and Hoechst (Thermo Fisher, Waltham, USA, H1399) for 10 min at RT, and mounted on slides using 40 µL of Vectashield. Images were captured using a Keyence BZ-X800 microscope and processed with BZ-X800 Analyzer software (Keyence, Osaka, Japan).

*Ex vivo imaging of mucosal surface*

Four weeks post transplantation, micro-optical coherence tomography (µOCT) imaging was performed as described previously (n=1 per group) [14]. Six measurements were taken at three regions of interest and averaged. Image analysis was conducted using ImageJ software (National Institutes of Health, Bethesda, USA, v1.51j8) to quantify airway surface liquid depth, periciliary liquid layer depth and mucociliary transport.

*Nasal potential difference (NPD) measurements*

NPD measurements were conducted to assess *in vivo* epithelial ion transport (n=3 per group) as described previously [14]. Polyethylene 90 tubing was positioned on the nasal septum for baseline readings, and on the bioscaffold site at day 28. Sequential perfusion of the following solutions was performed (≥ 5 min per solution): Ringer's lactate, Ringer's lactate with amiloride (100 µM), a low-chloride solution ($K_2HPO_4$ (2.4 mM), $K_H2PO_4$ (0.4 mM), Na Gluconate (115 mM), $NaHCO_3$ (25 mM), Ca $Gluconate_2$ (1.24 mM)), CFTR agonist (forskolin, 20 µM), and CFTR blockers (Inh-172, 10 µM + GlyH-101, 10 µM). The change in NPD (ΔNPD) following forskolin perfusion was calculated as: $NPD^{forskolin} – NPD^{low\ chloride}$.

*Statistical analysis*

Statistical guidance was provided by Stats Central, UNSW. All analyses were conducted using GraphPad Prism v9.0.1 (GraphPad Software, San Diego, USA). Data are presented as mean ± standard error of the mean (SEM). A *p*-value of <0.05 was considered statistically significant.

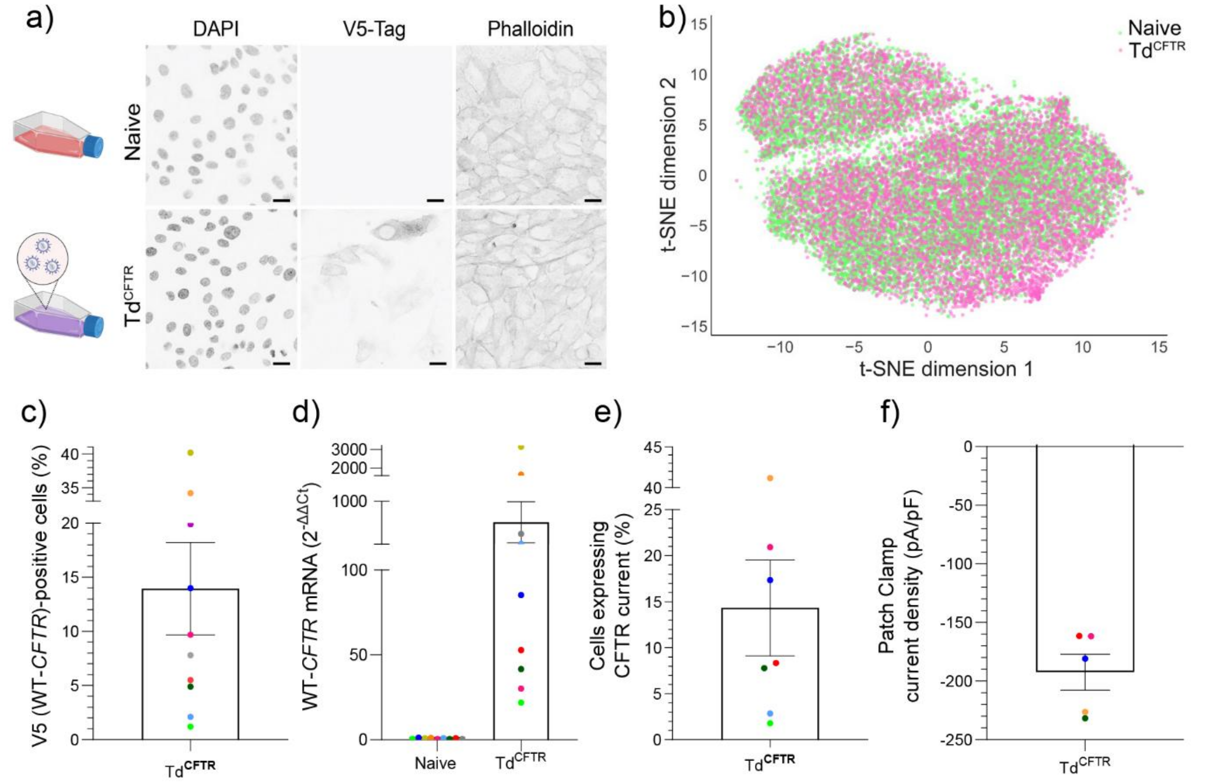


| PID | (%) | ($2^{-\Delta\Delta Ct}$) | (%) | (pA/pF) |
|---|---|---|---|---|
| 1 | 2.1 | 254.4 | 2.9 | n<5 |
| 2 | 4.9 | 41.7 | 7.8 | -231.8 ± 78.6 |
| 3 | 9.7 | 30.1 | 20.9 | -161.8 ± 33.1 |
| 4 | 40.2 | 3137.2 | ^ | ^ |
| 5 | 34.2 | 1664.5 | 41.2 | -226.4 ± 28.0 |
| 6 | 7.8 | 432.3 | ^ | ^ |
| 7 | 19.9 | ^ | ^ | ^ |
| 8 | 5.5 | 52.9 | 8.3 | -161.5 ± 42.3 |
| 9 | 14.0 | 85.2 | 17.4 | -180.9 ± 29.5 |
| 10 | 1.2 | 21.9 | 1.8 | n<5 |

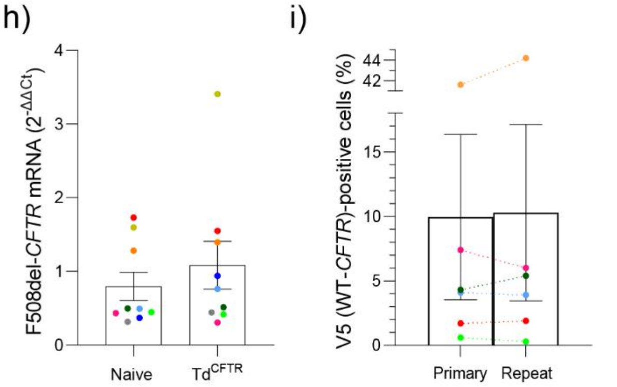


**Supplementary Figure 1. Lentiviral *CFTR* transduction of primary airway basal cells. a)** Representative confocal images of naive and transduced[CFTR] (Td$^{CFTR}$; MOI of 10) basal cell monolayers (PID 6) stained for nuclei (DAPI), the V5 epitope tag on the *CFTR* transgene, and actin (phalloidin). Inverted colour was used to enhance signal visibility. Scale bars: 10 µm. **b)** t-SNE plot of multivariate single-cell phenotypic data from naive and Td$^{CFTR}$ basal cells. **c)** Flow cytometric detection of V5 (*CFTR* transgene)-expressing cells (% of total Td$^{CFTR}$ basal cells). **d**) Relative *CFTR* transgene mRNA expression in Td$^{CFTR}$ basal cells ($2^{-\Delta\Delta Ct}$). **e**) Patch-clamp detection of Td$^{CFTR}$ basal cells expressing CFTR current (%). **f**) Patch-clamp current density measurements in Td$^{CFTR}$ basal cells (pA/pF). Data not shown for samples with n<5 basal cells expressing CFTR current. **g**) Data for flow cytometry (c), *CFTR* transgene mRNA expression (d), basal cells expressing CFTR current (e) and patch-clamp current density (f) in Td$^{CFTR}$ basal cells. ^Indicates no testing due to insufficient number of basal cells. **h)** Comparison of flow cytometry results following primary and repeat *CFTR* transduction. **i)** F508del

CFTR mRNA (ΔCt) expression in naive- and Td$^{CFTR}$-derived basal cells. Data are shown as mean ± SEM, with each dot representing an independent sample. PID=Participant ID. Statistical analysis was performed using paired t tests.

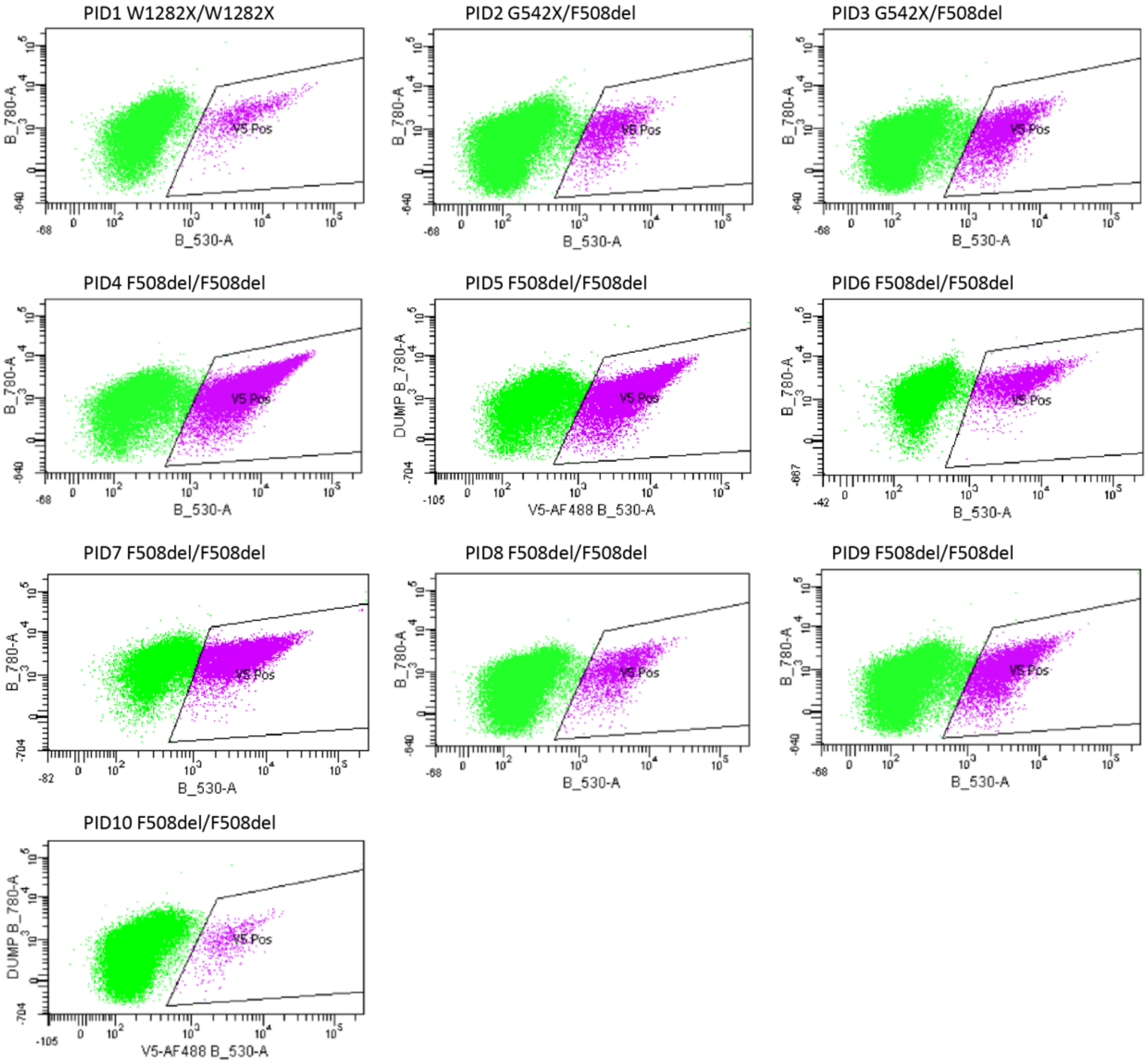


| | PID1 | | PID2 | | PID3 | | PID4 | | PID5 | |
|---|---|---|---|---|---|---|---|---|---|---|
| Population | # Events | % Total | # Events | % Total | # Events | % Total | # Events | % Total | # Events | % Total |
| All events | 50,000 | 100 | 50,000 | 100 | 50,000 | 100 | 50,000 | 100 | 50,000 | 100 |
| P1 | 25,777 | 51.6 | 38,883 | 77.8 | 41,135 | 82.3 | 37,143 | 74.3 | 32,775 | 65.6 |
| P2 | 24,184 | 48.4 | 38,469 | 76.9 | 40,949 | 81.9 | 36,881 | 73.8 | 32,658 | 65.3 |
| V5 pos | 1,048 | 2.1 | 2,456 | 4.9 | 4,862 | 9.7 | 20,079 | 40.2 | 17,094 | 34.2 |

| | PID6 | | PID7 | | PID8 | | PID9 | | PID10 | |
|---|---|---|---|---|---|---|---|---|---|---|
| Population | # Events | % Total | # Events | % Total | # Events | % Total | # Events | % Total | # Events | % Total |
| All events | 50,000 | 100 | 50,000 | 100 | 50,000 | 100 | 50,000 | 100 | 50,000 | 100 |
| P1 | 28,359 | 56.7 | 29,921 | 59.8 | 37,257 | 74.5 | 39,257 | 78.5 | 34,616 | 69.2 |
| P2 | 28,261 | 56.5 | 29,847 | 59.7 | 36,982 | 74 | 38,993 | 78 | 34,531 | 69.1 |
| V5 pos | 3,921 | 7.8 | 9,934 | 19.9 | 2,757 | 5.5 | 6,977 | 14 | 589 | 1.2 |

**Supplementary Figure 2. Participant-specific transduction efficiency at LV$^{CFTR}$ MOI 10.** Transduction efficiency was estimated by quantifying V5-positive cells via flow cytometry for each participant. PID=participant ID.

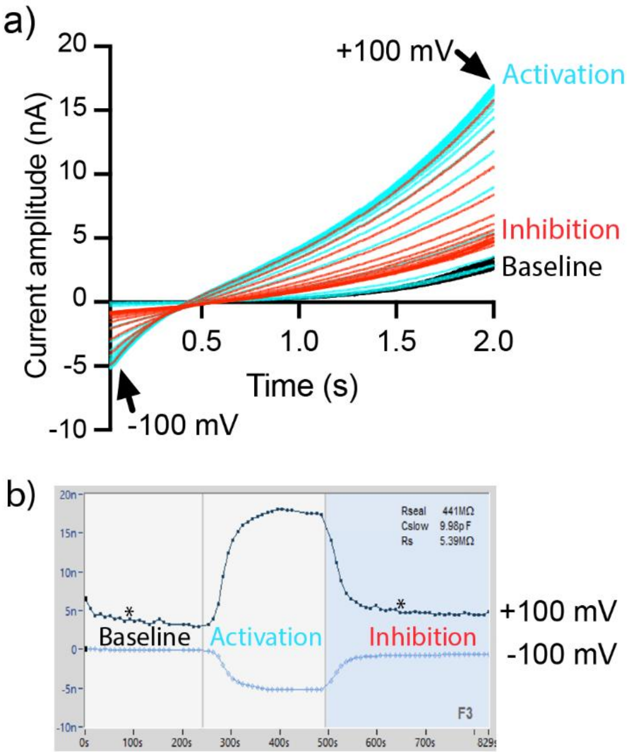


**Supplementary Figure 3. CFTR channel activity measured by whole cell patch-clamp electrophysiology. a)** Representative current traces showing baseline (black), CFTR activation (cyan) with 10 µM forskolin and 10 µM VX-770, and CFTR inhibition (red) with 10 µM CFTR-Inh$_{172}$. A ramp voltage protocol (-100 mV to +100 mV) was used to elicit CFTR currents. Black arrowheads indicate current amplitudes at +100 mV and -100 mV. Only the current traces between the two asterisks in (b) are shown. **b)** Time-lapse current amplitude measurements from (a), showing baseline, CFTR activation and CFTR inhibition at +100 mV and -100 mV.

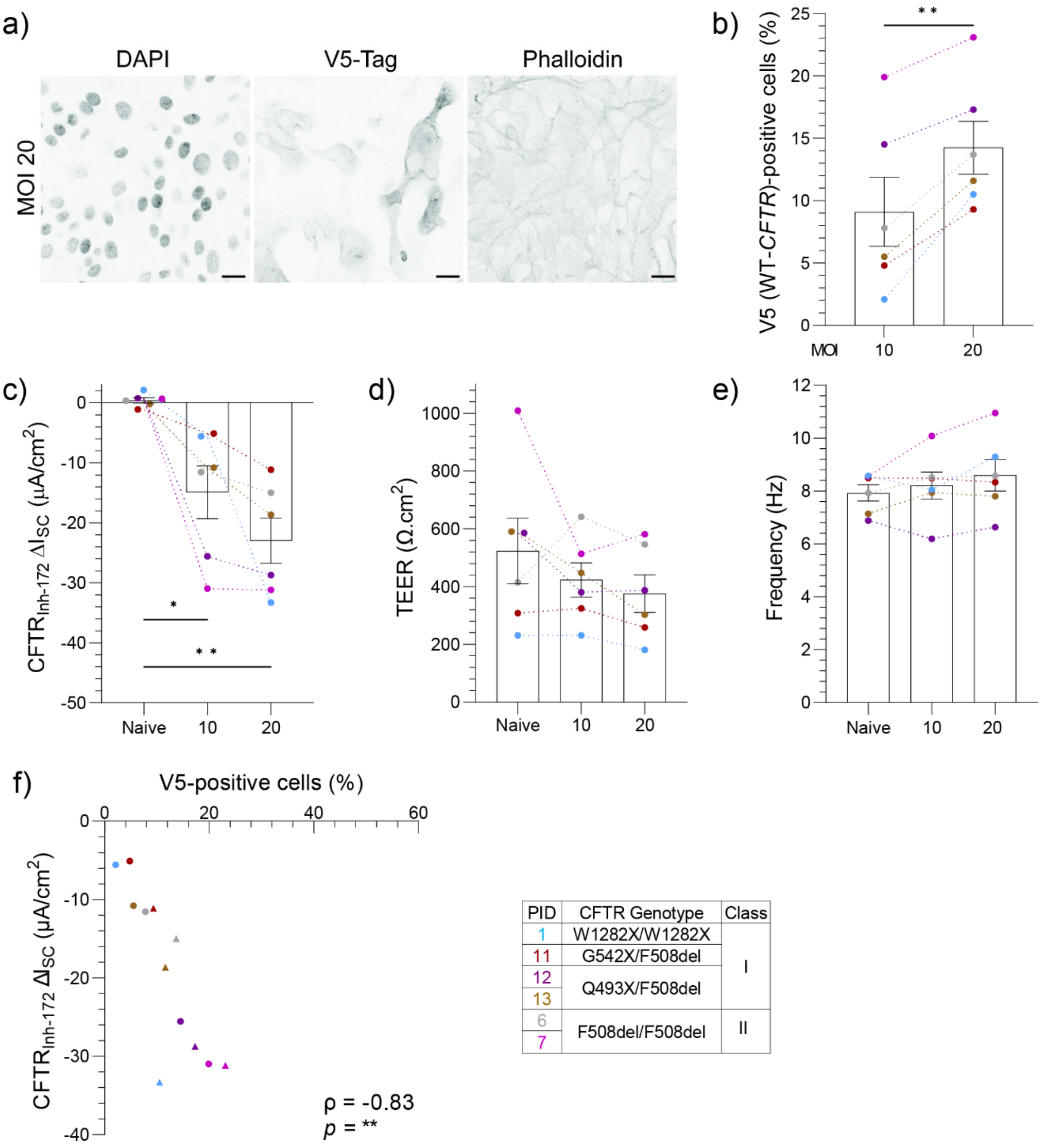


| PID | CFTR Genotype | Class |
|---|---|---|
| 1 | W1282X/W1282X | I |
| 11 | G542X/F508del | |
| 12 | Q493X/F508del | |
| 13 | | |
| 6 | F508del/F508del | II |
| 7 | | |

**Supplementary Figure 4. Transduction efficiency and functional assessment following LV*CFTR* transduction at MOI 10 versus MOI 20. a)** Representative confocal images of transduced[CFTR] (Td*CFTR*, MOI of 20) basal cell monolayers (PID 6) stained for nuclei (DAPI), the V5 epitope tag on the *CFTR* transgene, and actin (phalloidin). Inverted colour was used to enhance signal visibility. Scale bars: 10 μm. **b)** Flow cytometric detection of V5 (*CFTR* transgene)-expressing cells (% of total Td*CFTR* basal cells) at MOI of 10 and 20. **c-e)** Dot plots of **c)** mean CFTR-$Inh_{172}$ short-circuit current ($I_{SC}$), **d)** transepithelial electrical resistance (TEER; $\Omega.cm^2$) and **e)** cilia beat frequency (Hz) in naive and Td*CFTR*-derived (LV*CFTR* MOI of 10 and 20) differentiated epithelium of six CF participants (n=1 W1282X/W1282X, n=1 G542X/F508del, n=2 Q493X/F508del, n=2 F508del/F508del) with or without ETI treatment. **f)** Correlation between V5-positive cell percentage and CFTR-$Inh_{172}$ current ($\mu A/cm^2$)

in Td$^{CFTR}$-derived epithelium (MOI 10: circles, MOI 20: triangles). Each dot represents the average of n=3 replicate ALI cultures per participant. PID=Participant ID. ETI=elexacaftor/tezacaftor/ivacaftor. Statistical analysis was performed using a paired t test (a), One-way ANOVA with the Geisser-Greenhouse correction and Tukey's multiple comparison test (c, d, e) or Spearman's rank order correlation (d), *$p$<0.05, **$p$<0.01.

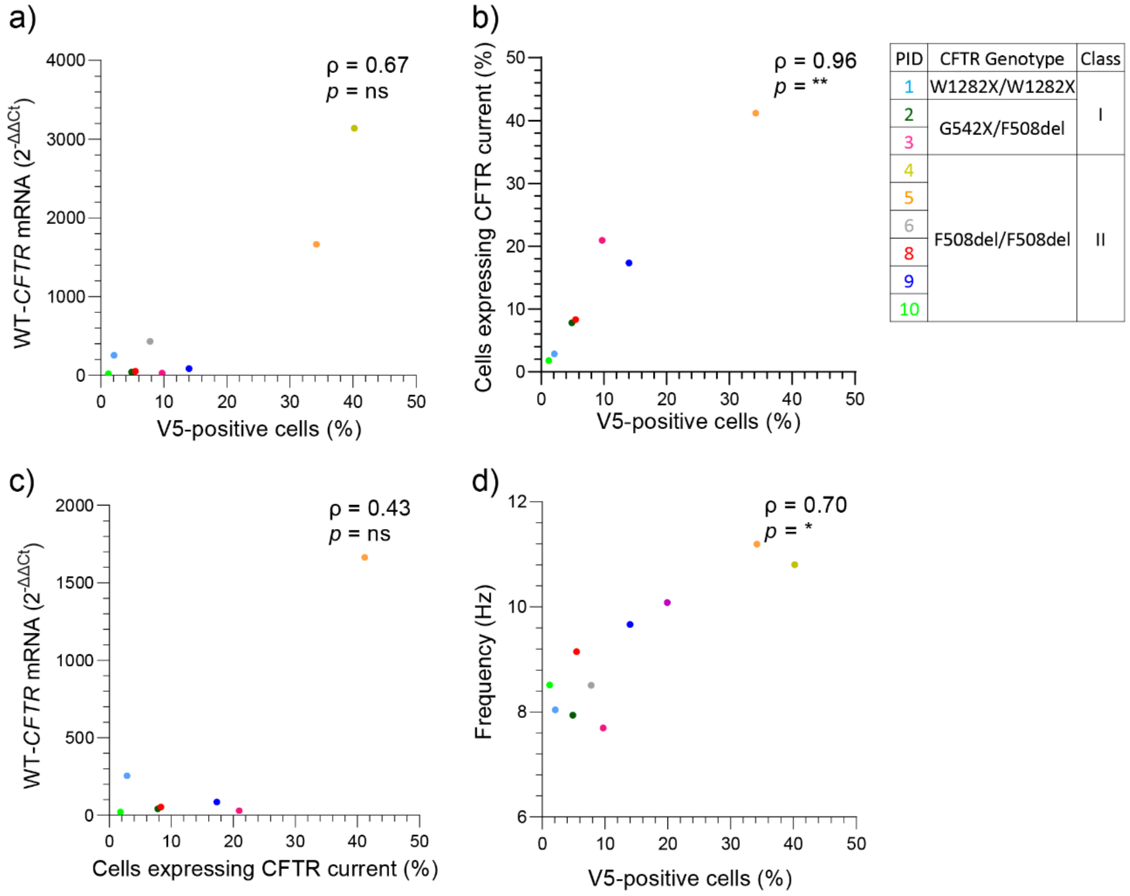


**Supplementary Figure 5. Correlation between *CFTR* expression and epithelial function metrics.** **a)** Correlation between V5 (*CFTR* transgene)-expressing cells (%) and *CFTR* transgene mRNA expression ($2^{-\Delta\Delta Ct}$). **b)** Correlation between cells expressing CFTR-Inh$_{172}$ current (%) and V5-positive cells (%). **c)** Correlation between CFTR-current expressing cells (%) and *CFTR* transgene mRNA expression ($2^{-\Delta\Delta Ct}$). **d)** Correlation between V5-positive cells (%) and cilia beat frequency (Hz). Data for PID 6 and PID 7 is not available due to the limited number of cells accessible for testing. Statistical analysis was performed using Spearman's rank order correlation. *$p$<0.05, **$p$<0.01. PID=Participant ID.

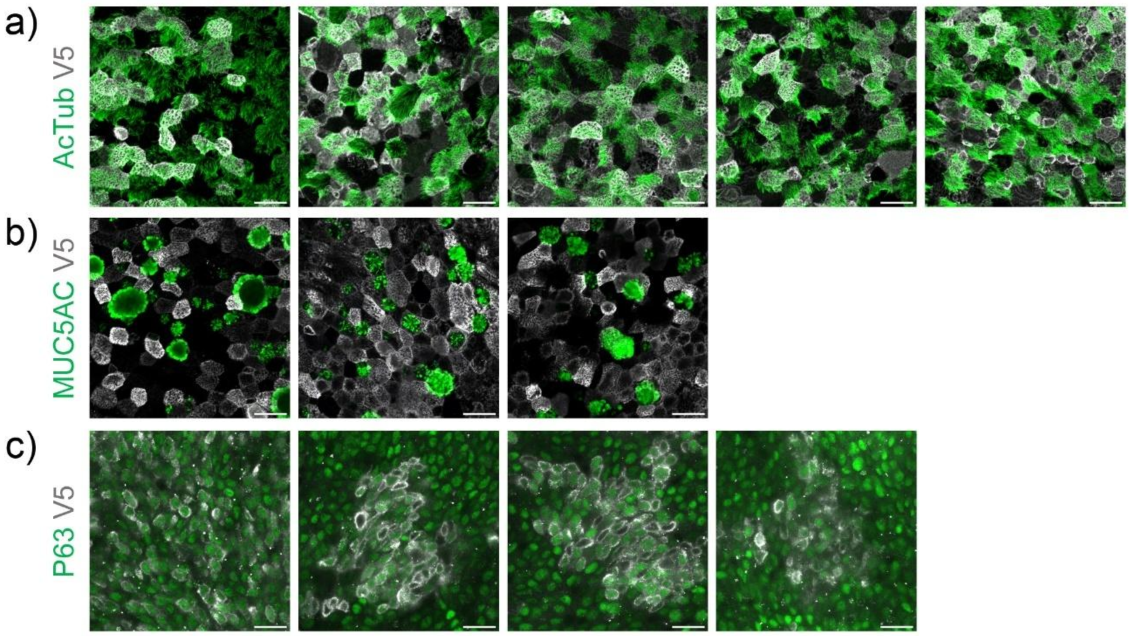


**Supplementary Figure 6. Immunofluorescence staining of transduced basal cells after air-liquid interface differentiation.** Replicate images showing co-location of the V5 epitope tag (grey) for CFTR with **a)** ciliated cells (acetylated tubulin, AcTub; green), **b)** secretory goblet cells (MUC5AC; green), and **c)** basal cells (p63; green). Images captured using a 63×/1.4 oil immersion objective. Scale bars = 20 μm.

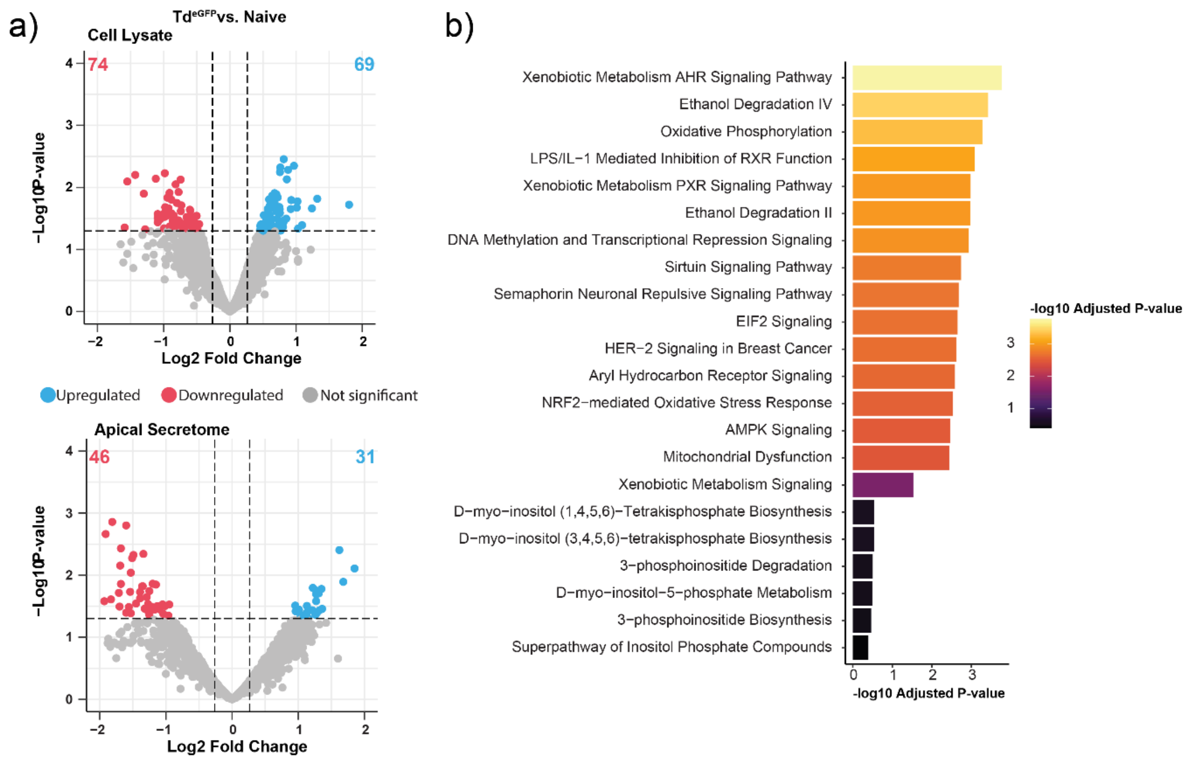


**Supplementary Figure 7. Proteomic profiling of eGFP-transduced CF airway basal cells differentiated at air-liquid interface. a)** Volcano plots of differentially abundant proteins in cell lysate (top, n=10) and apical secretome (bottom, n=3) from transduced ($Td^{eGFP}$)-derived epithelium relative to naive-derived epithelium. Each dot represents an individual protein, with upregulated proteins in blue and downregulated proteins in red. Coloured numbers indicate the total number of differentially abundant proteins. Dotted lines denote the significance cut-off ($p$-value $< 0.05$, fold change $> |1.2|$). Statistical analysis was performed using the DEP R Package (see methods). See Supplementary Table 3 for full data. **b)** Top enriched canonical pathways of differentially abundant cell lysate proteins between $Td^{eGFP}$-derived and naive-derived epithelium, identified using Ingenuity Pathway Analysis (IPA). Colour intensity represents the -$\log_{10}$ adjusted $p$-value.

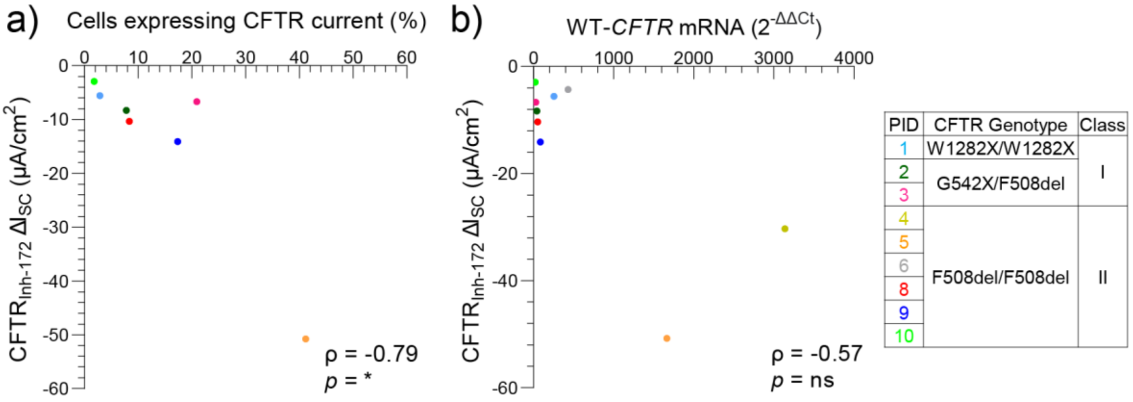


**Supplementary Figure 8. Correlation between *CFTR* expression and CFTR function metrics.** Correlation between $CFTR_{Inh\text{-}172}$ current ($\mu A/cm^2$) and **a)** cells expressing CFTR current (%) and **b)** *CFTR* transgene mRNA expression ($2^{-\Delta\Delta Ct}$). Statistical analysis was performed using Spearman's rank order correlation. **p*<0.05. PID=Participant ID.

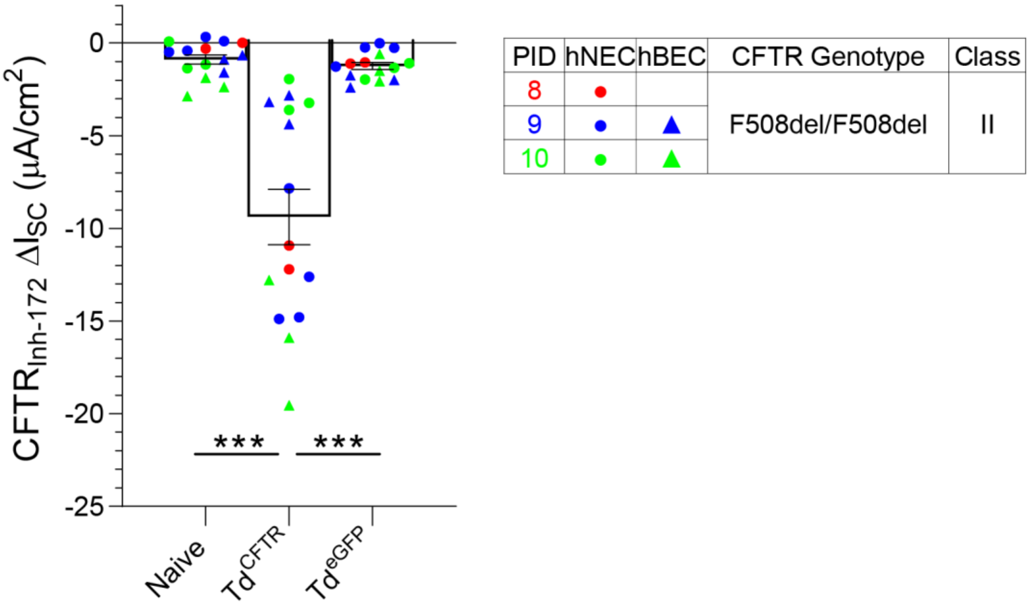


**Supplementary Figure 9. CFTR activity in transduced$^{eGFP}$-derived, transduced$^{CFTR}$-derived and naive-derived epithelium.** Dot plots of CFTR inhibited currents (CFTR-$Inh_{172}$) in naive or transduced ($Td^{CFTR}$ or $Td^{eGFP}$)-derived nasal (n=3) and bronchial (n=2) epithelium. Data are represented as mean ± SEM, with each dot representing the average of n=3 replicate ALI cultures per participant. PID=Participant ID. hNEC=human nasal epithelial cell. hBEC=human bronchial epithelial cell. Statistical analysis was performed using Brown-Forsythe and Welch ANOVA with Dunnett's T3 multiple comparisons test, ****p*<0.001.

a)

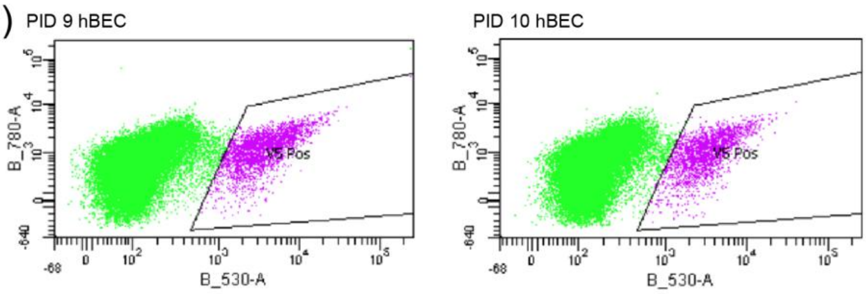


| | PID7 hBEC | | PID9 hBEC | | PID10 hBEC | |
|---|---|---|---|---|---|---|
| Population | # Events | % Total | # Events | % Total | # Events | % Total |
| All events | ^ | ^ | 50,000 | 100 | 50,000 | 100 |
| P1 | ^ | ^ | 38,851 | 77.7 | 35,487 | 71.0 |
| P2 | ^ | ^ | 38,695 | 77.4 | 35,259 | 70.5 |
| V5 pos | ^ | ^ | 2,658 | 5.3 | 2,340 | 4.7 |

b)

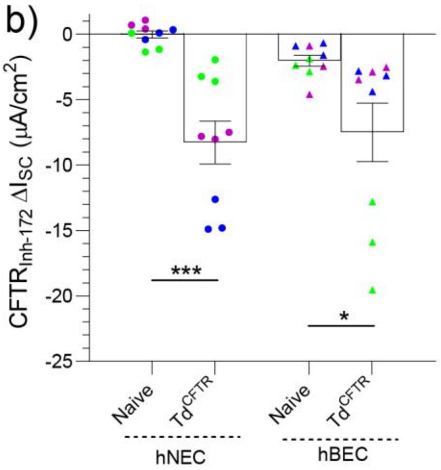


c)

| PID | V5-positive cells (%) | CFTR-Inh Isc (μA/cm²) |
|---|---|---|
| 7● | ^ | -7.8 ± 0.2 |
| 7▲ | ^ | -2.7 ± 0.2 |
| 9● | 14.0 | -14.1 ± 0.7 |
| 9▲ | 5.3 | -3.5 ± 0.5 |
| 10● | 1.2 | -2.9 ± 0.5 |
| 10▲ | 4.7 | -16.1 ± 2.0 |

● hNEC ▲ hBEC

**Supplementary Figure 10. Comparison of CFTR activity in nasal versus bronchial transduced-derived epithelium. a)** Human bronchial epithelial cell (hBEC) monolayer transduction efficiency (MOI of 10) estimated by quantifying V5-positive cells via flow cytometry. Refer to Supplementary Figure 2 for human nasal epithelial cell (hNEC) monolayer transduction efficiency. **b)** Dot plots of CFTR inhibited currents (CFTR-Inh$_{172}$) in naive- and transduced$^{CFTR}$ (Td$^{\mathit{CFTR}}$)-derived participant-matched hNEC and hBEC epithelium. **c)** Individual participant data for flow cytometry and CFTR-Inh$_{172}$. ^Indicates no testing due to insufficient number of basal cells. Data are shown as mean ± SEM, with each dot representing the average of n=3 replicate ALI cultures per participant. PID=Participant ID. Statistical analysis was performed using Welch's t test, *$p$<0.05, ***$p$<0.001.

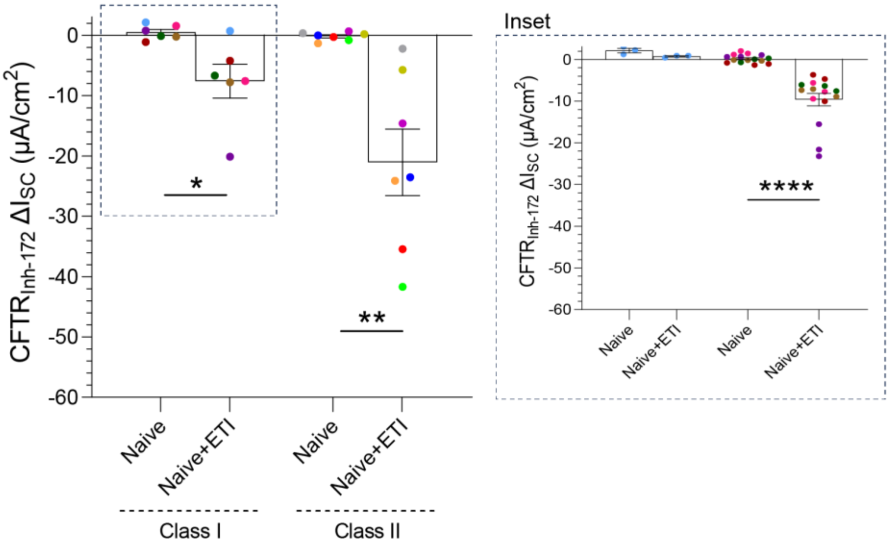


| PID | CFTR Genotype | Class |
|---|---|---|
| 1 | W1282X/W1282X | I |
| 2 | G542X/F508del | |
| 3 | | |
| 11 | | |
| 12 | Q493X/F508del | |
| 13 | | |
| 4 | F508del/F508del | II |
| 5 | | |
| 6 | | |
| 7 | | |
| 8 | | |
| 9 | | |
| 10 | | |

**Supplementary Figure 11. CFTR activity in ETI-treated epithelium.** Dot plots of CFTR inhibited currents in naive-derived and naive-derived ETI-treated epithelium from 10 CF participants with Class I (W1282X/W1282X (n=1), G542X/F508del (n=3), Q493X/F508del (n=2)) and Class II (F508del/F508del (n=7)) *CFTR* genotypes. Inset details Class 1 participants, showing n=3 replicate ALI cultures per participant. Data are shown as mean ± SEM, with each dot representing the average of n=3 replicate ALI cultures per participant. PID=Participant ID. ETI=elexacaftor/tezacaftor/ivacaftor. Statistical analysis was performed using a paired t test (a) or Welch's t test (b), *$p$<0.05, **$p$<0.01, ***$p$<0.0001.

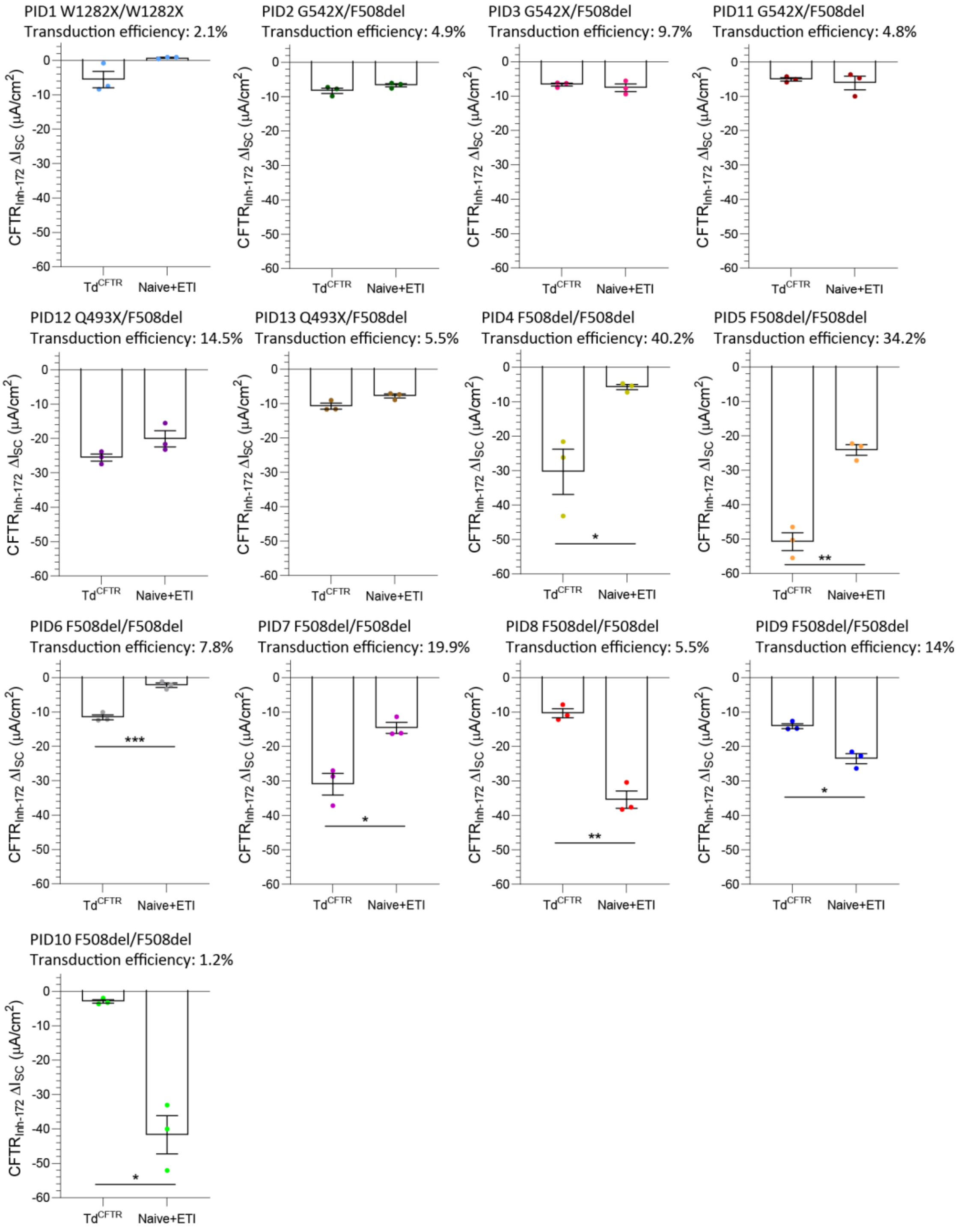


**Supplementary Figure 12. Individual participant data for CFTR activity across treatment groups.** Dot plots of CFTR-inhibited currents in transduced (Td$^{CFTR}$)-derived epithelium and naive-derived ETI-treated epithelium from 13 CF participants. Data from Figure 5 is presented per participant with transduction efficiency data from Supplementary Figure 2. Data are represented as Mean ± SEM, with each dot representing an independent ALI culture. PID=Participant ID.

ETI=elexacaftor/tezacaftor/ivacaftor. Statistical analysis was performed using Welch's t test, *$p$<0.05, **$p$<0.01, ***$p$<0.001.

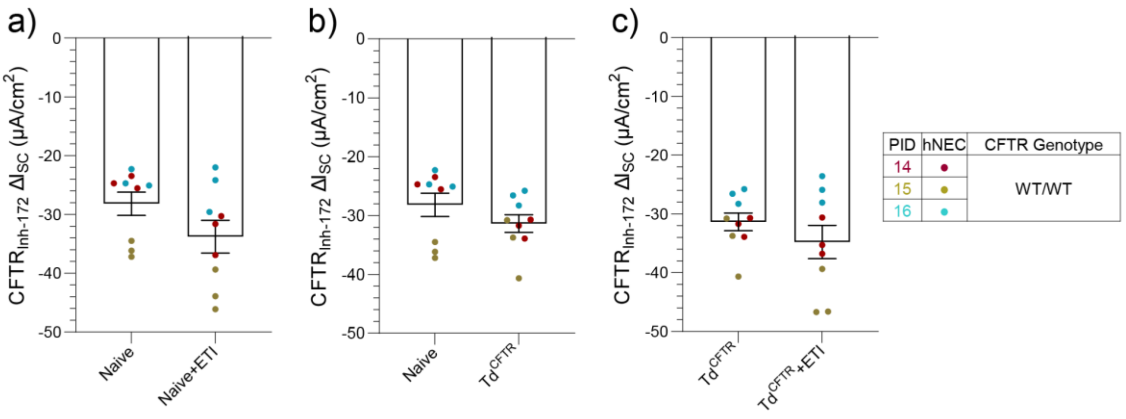


**Supplementary Figure 13. CFTR activity in non-CF epithelium following ETI, *CFTR* gene therapy, or their combination.** Dot plots of CFTR-inhibited currents (CFTR-Inh$_{172}$) in **a)** naive-derived and naive-derived ETI-treated epithelium, **b)** naive-derived and transduced$^{CFTR}$ (Td$^{CFTR}$)-derived epithelium and **c)** Td$^{CFTR}$-derived and Td$^{CFTR}$-derived ETI-treated epithelium from three non-CF participants. Data are shown as mean ± SEM, with each dot representing an independent ALI culture. PID=Participant ID. ETI=elexacaftor/tezacaftor/ivacaftor. Statistical analysis was performed using Welch's t test.

**Supplementary Table 1. Demographics of study participants.**

| PID | CFTR functional defect | CFTR genotype | Age (years) | Transduced | |
|---|---|---|---|---|---|
| | | | | CFTR | eGFP |
| 1 | Class I – Protein production defect | W1282X/W1282X | 11.5 | ✓ | |
| 2 | | G542X/F508del | 12.7 | ✓ | |
| 3 | | G542X/F508del | 2.0 | ✓ | |
| 11 | | G542X/F508del | 8.51 | ✓ | |
| 12 | | Q493X/F508del | 4.15 | ✓ | |
| 13 | | Q493X/F508del | 7.21 | ✓ | |
| 4 | Class II – Folding/maturation defect | F508del/F508del | 14.7 | ✓ | |
| 5 | | F508del/F508del | 14.9 | ✓ | |
| 6 | | F508del/F508del | 13.4 | ✓ | |
| 7 | | F508del/F508del | 1.7 | ✓ | |
| 8 | | F508del/F508del | 11.2 | ✓ | ✓ |
| 9 | | F508del/F508del | 4.8 | ✓ | ✓ |
| 10 | | F508del/F508del | 4.0 | ✓ | ✓ |
| 14 | non-CF | WT/WT | 0.9 | ✓ | |
| 15 | | | 5.7 | ✓ | |
| 16 | | | 1.2 | ✓ | |
| 17 | | | 14.4 | | |
| 18 | | | 4.5 | | |
| 19 | | | 3.8 | | |
| 20 | | | 1.4 | | |
| 21 | | | 2.5 | | |
| 22 | | | 11.9 | | |
| 23 | | | 7.7 | | |

PID=Participant ID; WT=Wild-Type; eGFP=enhanced Green Fluorescent Protein.

**Supplementary Table 2. Antibodies used for immunofluorescence staining.**

| Antibody | Target | Supplier | Catalogue Number | Dilution |
|---|---|---|---|---|
| Anti-p63 antibody [4A4] | Basal cell | Abcam | ab735 | 1:250 |
| Monoclonal anti-Acetylated Tubulin | Ciliated cell | Sigma-Aldrich | T7451 | 1:250 |
| Monoclonal anti-MUC5AC (45M1) | Goblet cell | Thermo Fisher Scientific | MA5-12178 | 1:250 |
| V5-Tag (D3H8Q) Rabbit mAb | CFTR | Cell Signaling Technology | #13202 | 1:50 |
| Phalloidin-Atto 565 | Actin | Sigma-Aldrich | 94072 | 1:2000 |
| DAPI (4',6-Diamidino-2-Phenylindole, Dihydrochloride) | DNA | Thermo Fisher Scientific | D1306 | 1:5000 |
| Alexa Fluor 488 goat anti-rabbit IgG | | Thermo Fisher Scientific | A-11034 | 1:500 |
| Alexa Fluor 647 goat anti-mouse IgG | | Thermo Fisher Scientific | A-21236 | 1:500 |

**Supplementary Table 3. Differentially abundant proteins in cell lysate and apical secretome of naive-derived versus LV$^{CFTR}$ and LV$^{eGFP}$ transduced-derived epithelium.**

Provided as supplemental spreadsheet.

**Supplementary Table 4. Data for the short-circuit currents and electrophysiological parameters in naive, LV$^{CFTR}$-transduced and LV$^{eGFP}$-transduced derived epithelium (n=13 CF; n=10 non-CF).**

Provided as supplemental spreadsheet. Data represent transepithelial resistance, short-circuit current values for forskolin stimulated cAMP currents (ΔFsk) and CFTR-Inh$_{172}$ inhibited currents (ΔFsk + CFTR-Inh$_{172}$). Values represented are after nil or ETI treatment (VX-445 (3µM/48h) + VX-661 (18µM/48h) + VX-770 (10µM/acute)). Mean (±SEM); PID = Participant ID.

**Supplementary Table 5. *CFTR* genotype-specific data for naive and LV$^{CFTR}$ transduced (Td$^{CFTR}$)-derived epithelium in the presence and absence of ETI.**

| *CFTR* Genotype | Mean $I_{SC}$ CFTR-$Inh_{172}$ | | | |
|---|---|---|---|---|
| | Naive | $Td^{CFTR}$ | Naive+ETI | $Td^{CFTR+}$ETI |
| W1282X/W1282X (n=1) | 2.13 ± 0.52 | -5.57 ± 2.38 | 0.73 ± 0.17 | -5.32 ± 1.56 |
| G542X/F508del (n=2) Q493X/F508del (n=3) | 0.17 ± 0.45 | -11.28 ± 0.99 | -9.67 ± 1.51 | -26.12 ± 2.91 |
| F508del/F508del (n=7) | -0.21 ± 0.25 | -17.22 ± 6.57 | -21.49 ± 5.19 | -37.81 ± 4.98 |
| non-CF (n=3) | -28.18 ± 1.98 | -31.36 ± 1.50 | -33.77 ± 2.79 | -34.78 ± 2.82 |

**Supplementary Video 1. Video showing placement of a cell-free scaffold onto the rabbit nasal septum.**

Provided as video file.

**Supplementary Video 2. Video showing placement of a scaffold+cell graft onto the rabbit nasal septum.**

Provided as video file.

**References**

1. Hewson CK, Capraro A, Wong SL, et al. Novel Antioxidant Therapy with the Immediate Precursor to Glutathione, gamma-Glutamylcysteine (GGC), Ameliorates LPS-Induced Cellular Stress in In Vitro 3D-Differentiated Airway Model from Primary Cystic Fibrosis Human Bronchial Cells. Antioxidants (Basel). 2020;9(12).
2. Allan KM, Wong SL, Fawcett LK, et al. Collection, Expansion, and Differentiation of Primary Human Nasal Epithelial Cell Models for Quantification of Cilia Beat Frequency. J Vis Exp. 2021(177):e63090.
3. Awatade NT, Wong SL, Capraro A, et al. Significant functional differences in differentiated Conditionally Reprogrammed (CRC)- and Feeder-free Dual SMAD inhibited-expanded human nasal epithelial cells. J Cyst Fibros. 2021;20(2):364-71.
4. Carpenter AE, Jones TR, Lamprecht MR, et al. CellProfiler: image analysis software for identifying and quantifying cell phenotypes. Genome Biol. 2006;7(10):R100.
5. Van der Maaten L, Hinton G. Visualizing data using t-SNE. Journal of machine learning research. 2008;9(11).
6. Allan KM, Astore MA, Kardia E, et al. Q1291H-CFTR molecular dynamics simulations and ex vivo theratyping in nasal epithelial models and clinical response to elexacaftor/tezacaftor/ivacaftor in a Q1291H/F508del patient. Front Mol Biosci. 2023;10:1148501.

7. Clarke LA, Awatade NT, Felicio VM, et al. The effect of premature termination codon mutations on CFTR mRNA abundance in human nasal epithelium and intestinal organoids: a basis for read-through therapies in cystic fibrosis. Hum Mutat. 2019;40(3):326-34.
8. Perez-Riverol Y, Bai J, Bandla C, et al. The PRIDE database resources in 2022: a hub for mass spectrometry-based proteomics evidences. Nucleic Acids Res. 2022;50(D1):D543-D52.
9. Cox J, Neuhauser N, Michalski A, et al. Andromeda: a peptide search engine integrated into the MaxQuant environment. J Proteome Res. 2011;10(4):1794-805.
10. Cox J, Hein MY, Luber CA, et al. Accurate proteome-wide label-free quantification by delayed normalization and maximal peptide ratio extraction, termed MaxLFQ. Mol Cell Proteomics. 2014;13(9):2513-26.
11. Zhang X, Smits AH, van Tilburg GB, et al. Proteome-wide identification of ubiquitin interactions using UbIA-MS. Nat Protoc. 2018;13(3):530-50.
12. Kramer A, Green J, Pollard J, Jr., et al. Causal analysis approaches in Ingenuity Pathway Analysis. Bioinformatics. 2014;30(4):523-30.
13. Wong SL, Awatade NT, Astore MA, et al. Molecular Dynamics and Theratyping in Airway and Gut Organoids Reveal R352Q-CFTR Conductance Defect. Am J Respir Cell Mol Biol. 2022;67(1):99-111.
14. Cho DY, Zhang S, Skinner DF, et al. Ivacaftor restores delayed mucociliary transport caused by Pseudomonas aeruginosa-induced acquired cystic fibrosis transmembrane conductance regulator dysfunction in rabbit nasal epithelia. Int Forum Allergy Rhinol. 2022;12(5):690-8.
15. Bohn GA, Chaffin AE. Extracellular matrix graft for reconstruction over exposed structures: a pilot case series. J Wound Care. 2020;29(12):742-9.
16. Bosque BA, Dowling SG, May BCH, et al. Ovine Forestomach Matrix in the Surgical Management of Complex Lower-Extremity Soft-Tissue Defects. J Am Podiatr Med Assoc. 2023;113(3).
